# Dihydromyricetin promotes GLP-1 secretion to improve insulin resistance via “gut microbiota-CDCA”

**DOI:** 10.1101/2024.05.28.596357

**Authors:** Pengfei Li, Yong Zhang, Hedong Lang, Pengfei Hou, Yu Yao, Ruiliang Zhang, Xiaolan Wang, QianYong Zhang, Mantian Mi, Long Yi

**Affiliations:** Research Center for Nutrition and Food Safety, Chongqing Key Laboratory of Nutrition and Health, Chongqing Medical Nutrition Research Center, Institute of Military Preventive Medicine, Third Military Medical University, Chongqing 400038, P.R. China.

**Keywords:** Chenodeoxycholic acid, Dihydromyricetin, Glucagon-like peptide-1, Gut microbiota, Insulin resistance

## Abstract

Dihydromyricetin (DHM) is a polyphenolic phytochemical found mainly in plants such as *Ampelopsis grossedentata,* which has beneficial effects on insulin resistance. However, the specific mechanism has not been clarified. In this study, C57BL/6 mice were exposed to a high-fat diet (HFD) for eight weeks. DHM could improve insulin resistance via enhancing the incretin effect. DHM increased serum GLP-1 by improving intestinal GLP-1 secretion and inhibiting GLP-1 decomposition, associated with the alteration of intestinal intraepithelial lymphocytes (IELs) proportions and decreased expression of CD26 in IELs and TCRαβ^+^ CD8αβ^+^ IELs in HFD-induced mice. Meanwhile, DHM could ameliorate GLP-1 level and insulin resistance by modulation of gut microbiota and the metabolites, particularly the regulation of intestinal bile acid CDCA content, followed by the inhibition of FXR expression in intestinal L cells as well as increased Gcg mRNA expression and the secretion of GLP-1. These findings clarify the role of the “gut microbiota-CDCA” pathway in the improvement of intestinal GLP-1 levels in HFD-induced mice by DHM administration, providing a new pharmacological target for the prevention of insulin resistance.

## 1. Introduction

Insulin resistance was defined as the inability of insulin to optimally stimulate glucose transport to body cells (hyperinsulinemia or impaired glucose tolerance)(Lee *et al*, 2022). Over the past decades, it has become clear that insulin resistance is a common risk factor for many diseases, including diabetes mellitus type 2 (T2DM), cardiovascular diseases and endocrine diseases (Bovolini *et al*, 2021; Piché *et al*, 2020). There are currently no drugs specifically approved for the treatment of insulin resistance, but various studies have demonstrated the efficacy of certain antidiabetic drugs in improving insulin resistance (Lee *et al*, 2022), such as GLP-1 receptor agonists (GLP-1RA)(Marušić *et al*, 2021, 2). GLP-1 is an incretin secreted by L cells of intestinal mucosa (Drucker, 2022). It has the function of enhancing glucose-dependent insulin secretion, inhibiting glucagon secretion, slowing gastric emptying to reduce blood glucose and alleviating insulin resistance(Müller *et al*, 2019; Smith *et al*, 2019). GLP-1 is rapidly degraded by intestinal CD26/DPP-IV (dipeptidyl peptidase IV) after production (Müller *et al*, 2019). Dipeptidyl peptidase 4 (DPP-4/CD26), a serine protease belonging to the type II transmembrane glycoprotein family, is expressed on the surface of T cells, B cells, and myeloid cells (Aliyari Serej *et al*, 2017). Studies show that serum GLP-1 level is significantly lower in patients with insulin resistance, so maintaining the level of GLP-1 is of great significance to prevent or delay the development of insulin resistance (Drucker, 2022). Although GLP-1 receptor agonists have achieved some efficacy in the treatment of insulin resistance, there are also some drawbacks, such as side effects, poor compliance and drug resistance (Wilding *et al*, 2021; Shetty *et al*, 2022). Thereby, approaches aimed at boosting the endogenous production or release of GLP-1 is regarded as an innovative approach for treating insulin resistance.

Dihydromyricetin (DHM) is a polyphenolic phytochemical found in plants such as *Ampelopsis grossedentata* (Zhang *et al*, 2018). Our previous studies found that regular DHM supplementation could significantly alleviate the metabolic abnormalities of fasting blood glucose, glycosylated hemoglobin, cystatin C in adults with type 2 diabetes (Ran *et al*, 2019). DHM notably promoted the maintenance of intestinal mucosal barrier integrity through regulating gut microbiota and short-chain fatty acids in nonalcoholic steatohepatitis (NASH) C57BL/6 mice model induced by high fat diet (HFD). In addition, DHM significantly reversed the elevated blood glucose abnormally in HFD-induced mice (Wang *et al*, 2022). DHM could also increase serum GLP-1 level and enhance exercise endurance in aerobic exercise mice (L *et al*, 2022), indicating that DHM may play a role in preventing and treating chronic diseases by regulating gut microbiota or its metabolites to increase GLP-1 level in vivo. Published literatures indicated that DHM combined with *L. rhamnosus* 1.0320 restored the intestinal barrier induced by acute alcohol exposure by upregulating the intestinal short-chain fatty acid levels (Wang et al, 2023a). However, the underlying molecular mechanism involved with the enhancement of GLP-1 by DHM administration remains to be further elucidated.

Recent studies have found that intestinal regional immunity is closely related to the development of insulin resistance, obesity and other diseases. Intestinal intraepithelial lymphocytes (IELs) are a unique population of T cells located in the intestinal mucosa (Olivares-Villagómez & Kaer, 2018; Glaubitz *et al*, 2023). Its different subtypes and functional disorders play an important role in the occurrence of various metabolic diseases (Bordon, 2019; He *et al*, 2019). It has been reported that integrin β7^+^ IELs are involved in the development of insulin resistance via regulating the frequency of L cells and level of GLP-1 (He *et al*, 2019). The result suggested that some special subsets of intestinal IELs may affect the expression level of gut-derived GLP-1 and regulate the metabolism of the body. Our previous work showed that DHM increased the proportion of ILC3 cells and IL-22 secreted by ILC3 cells in colonic lamina propria, and promoted the expression level of IL-22, implying a potential role of DHM on the modulation of intestinal regional immunity (Zhou *et al*, 2023). In this study, we hypothesized that DHM could promote GLP-1 level and improve insulin resistance in HFD-induced mice by modulation of gut microbiota and the metabolites, which might be associated with modulation of the intestinal L cells and certain subsets of IELs cells.

## 2. Materials and methods

### 2.1 Animals and Experimental Protocol

Animal experiments were approved by the ethics committee of the Army Medical University (Chongqing, China) and performed following the guidelines of the center of laboratory animals at the Army Medical University. C57BL/6 mice (SPF grade, male, 6-8 weeks old) were purchased from Hunan SJA Laboratory Animal Co. Ltd. (Hunan, China) and allowed one week of acclimatization. The mice were maintained under specific pathogen free conditions in a controlled facility with free access to chow and water (temperature 20-22 °C, humidity 45 ±5%, 12 h light/dark cycle).

Mice were randomly divided into three groups with different diets administration for 8 weeks: (1) Control group (n=10): chow diet (10% kcal from fat, 70% kcal from carbohydrate, 20% kcal from protein); (2) HFD group (n=10): High-fat diet (45% kcal from fat, 35% kcal from carbohydrate, 20% kcal from protein); (3) HFD with DHM group (n=10): HFD containing 0.6% DHM (equivalent to 300 mg/kg·B.W./day). Body weight and food intake were measured once a week during the experiments. Average daily food intake (g/day/mouse) was calculated. Feces samples were collected at the 8th week for 16S rRNA, untargeted metabolomics and bile acid (BA) assessment. Mice were fasted overnight and then anesthetized for harvesting blood and ipeptidyl peptidase-4 (DPP4) enzyme inhibitor (final concentration of 5 μM) was added. The blood was centrifuged at 3000×g for 10 min for serum isolation. Intestinal tissues were collected immediately after euthanasia. All samples were stored at −80 °C until used for analysis. And the intestinal IELs were isolated by flow cytometry. For gut microbiota clearance experiment, after HFD with or without DHM administration for 4 weeks, mice were fed with water containing antibiotics in the last 4 weeks. For gut microbiota transplantation experiment, mice were administrated with HFD with or without DHM for 8 weeks, and were fed with antibiotics at the 5th and 6th week, followed by the stomach gavage of feces at the 7th and 8th week. All animal procedures were supervised and approved by the animal care committee at the Third Military Medical University (Chongqing, China; Approval AMUWEC20211008).

### 2.2 Biochemical analysis

Serum glucose, TG, LDL-C, HDL-C and CHO were measured using an automatic biochemical analyser. Serum GLP-1 were measured using a commercial biochemical kit (Jingmei, China).

### 2.3 Oral glucose tolerance test (OGTT) and intraperitoneal glucose tolerance test (IPGTT)

After fasting overnight, the basal blood glucose level was measured first, and mice were given oral glucose or intraperitoneal glucose (2 g/kg), and the blood glucose levels were measured with a blood glucose meter at 0, 15, 30, 60, 90, and 120 min. Glucose tolerance impairment was assessed by area under the curve (AUC).

### 2.4 Insulin resistance test (ITT)

During the insulin tolerance test, the mice were intraperitoneally injected with insulin (0.75 U/kg) after fasting for 6 h, and the blood glucose level were measured by blood glucose meter at 15, 30, 60, 90 and 120 min. Insulin resistance was assessed by the area under the curve (AUC).

### 2.5 Immunofluorescence staining

For immunofluorescence staining of intestine tissues (n=3 per group), the fresh intestinal tissues were embedded in optimal cutting temperature (OCT) compound and cut into frozen intestinal sections. The slides were treated with GLP-1 antibody, and were then exposed to Alexa Fluor 488 goat Anti-rabbit IgG and Alexa Fluor 594 goat anti-rabbit IgG. Nuclei were stained with 4,6-diamidino-2-phenylindole-dihydrochloride (DAPI). After PBS washing, slides were mounted using Prolong^Tm^ Gold Antifade Mountant (Life Technologies) and were photographed with a fluorescence microscope camera.

### 2.6 DNA extraction, 16S rRNA and Illumina MiSeq sequencing

Microbial genomic DNA were extracted from each stool sample Using E.Z.N.A.®Soil DNA Kit (Omega Bio-tek, USA). The extracted DNA was detected on a 1% agarose (biowest, ES) gel, and the concentration and purity of DNA were determined using a NanoDrop 2000 ultra-micro spectrophotometer (Thermo Fisher Scientific, USA). The V3-V4 hypervariable region of the bacterial 16S rRNA gene was amplified using primer pairs. 20 μL PCR reaction system include 1 μL (10 ng) template DNA, 0.8 μL (5 μM) forward and reverse primers each, 0.4 μL FastPfu DNA Polymerase, 4 μL 5×FastPfu Buffer and 2 μL (2.5 mM) dNTPs. The PCR reaction conditions were listed as follows: initial denaturation at 95℃ for 30 min, followed by 30 cycles of denaturation at 95℃ for 30 s, annealing at 55℃ for 30 s, extension at 72℃ for 45 s, and stable extension at 72℃ for 10 min until that end of the reaction. The PCR products were examined by gel electrophoresis on a 2% agarose gel, and then the PCR products were purified using AxyPrep DNA Gel Extraction Kit (Axygen Biosciences, Axygen, USA) according to the manufacturer’s instructions. PCR products were quantified using a Quantus™ Fluorometer (Promega, USA). According to the sequencing requirements of each sample, the corresponding proportion of mixing was carried out. Use the NEXTFLEX Rapid DNA-Seq Kit to build the database. The NovaSeq PE250 platform of Illumina was used for sequencing. The sequences were OTU clustered according to 97% similarity using the UPARSE software (http://drive5.com/uparse/). All data analysis was carried out on the Meiji Biological Cloud Platform (https://cloud.majorbio.com).

### 2.7 Untargeted metabolomics analysis

A 50 mg fecal sample was placed in a 2 mL of centrifuge tube with a 6 mm diameter ground bead. Metabolites were extracted with 400 μL of extract (methanol: water = 4:1 (V: V)) containing 0.02 mg/mL internal standard (L-2-chlorophenylalanine). The sample solution was triturated for 6 min (−10℃, 50 Hz) in the cryo-tissue triturator, and then extracted with low temperature ultrasound for 30 min (5℃, 40 kHz). The sample was kept at −20℃ for 30 min, centrifuged for 15 min (4℃, 13000 ×g), and the supernatant was transferred to a sampling vial with an inner cannula for analysis. Quality control (QC) samples were prepared by mixing the metabolites of all samples in the same volume. During the instrumental analysis, one QC sample was inserted into every four samples to examine the repeatability of the whole analysis process. The instrument platform of this LC-MS analysis is the UHPLC-Q Exactive HF-X system of Thermo Flying Company. After the operation, the LC-MS raw data was imported into the metabonomics processing software Progenesis QI (Waters Corporation, Milford, USA) for baseline filtering, peak identification, integration, retention time correction, and peak alignment. Finally, a data matrix of retention time, mass-to-charge ratio and peak intensity is obtained. At the same time, the MS and MSMS mass spectra were matched with the public metabolic databases HMDB (http://www.hmdb.ca/) and Metlin (https://metlin.scripps.edu/), as well as the Meiji self-built database to obtain metabolite information. After searching the database, the matrix data is uploaded to the Meiji Biological Cloud Platform (https://cloud.majorbio.com) for data analysis. The metabolic pathway annotation of the differential metabolites was performed through the KEGG database (https://www.kegg.jp/kegg/pathway.html), and the pathways involved in the differential metabolites were obtained. Pathway enrichment analysis was performed by the Python package of scipy.stats, and the biological pathways most relevant to the experimental treatment were obtained by Fisher’s exact test.

### 2.8 Bile Acids (BAs) targeted metabolomics analysis

Precisely pipette 100 μL of sample, add 50 μL of internal standard working solution (200 ng/mL), then add 350 μL of extraction solution (methanol), vortex and mix for 30 s, perform low temperature ultrasound for 30 min (5℃, 40 KHz), stand for 30 min at −20℃, and centrifuge for 15 min at 4℃, 13000 ×g. The supernatant was dried with nitrogen, redissolved with 100 μL of 50% acetonitrile water, vortexed and mixed for 30 s, ultrasonically treated at low temperature for 10 min (5℃, 40 KHz), centrifuged at 4℃ for 15 min with 13000 ×g, and the supernatant was detected by LC-MS/MS. The samples were analyzed by LC-MS/MS using an Exion LC AD liquid phase system in combination with a QTRAP® 6500+ mass spectrometer (Shanghai Meiji Biomedical Technology Co., Ltd.).

After the operation, the original data of LC-MS was imported into Sciex quantitative software OS for automatic identification and integration of each ion fragment with default parameters, and manual inspection was assisted to draw a linear regression standard curve with the ratio of the peak area of the analyte to the peak area of the internal standard as the ordinate and the concentration of the analyte as the abscissa. Sample concentration calculation: substitute the ratio of peak area of sample analyte to peak area of internal standard into the linear equation to calculate the concentration result.

### 2.9 IELs isolation and flow cytometry

After mice were sacrificed, the small intestinal tissues in each group were quickly separated, and the mesentery and adipose tissues were removed in ice-cold PBS. The intestinal tissues were cut open along the longitudinal axis, and the contents were removed and washed twice in ice-cold PBS. Small intestinal tissue was cut into small pieces and put into 50 mL EP tubes, each tube was added with 20 mL of digestive juice (5% FBS+D-Hanks+1 mM DTT+5 mM EDTA+10 mM HEPES), and then shaken for 20 min on a constant temperature shaker at 37 ℃ and 210 × rpm. After shaking, swirl the EP tube containing the digestion solution for 20 s, filter and collect the digestion solution into a new 50 mL EP tube using a 70 μm strainer, and add 20 ml to the tube, repeat the above steps, and collect the filtrate. The pooled digest was centrifuged at 2000 × rpm for 5 min, and the supernatant was discarded. The pellet was resuspended in 9 mL of 40% Percoll and transferred to a 15 mL round-bottomed EP tube. Add 5 mL of 70% Percoll to the bottom of each EP tube through a pipette, centrifuge at 900 × g at 20℃, horizontal density gradient for 20 min, and adjust the acceleration and deceleration of the centrifuge to 0. After centrifugation, the round-bottomed EP tube was carefully collected to the pipette rack. The liquid was divided into three layers, and the middle white layer was the IELs cells. Use a Buss pipette to suck 4mL of the intermediate tunica albuginea into a 15ml EP tube with a sharp bottom, and add 8mL of washing solution (PBS+5%FBS) for washing, then centrifuge at 2000 × rpm for 5 min, and collect the precipitate as IELs.

### 2.10 Cell culture and enzyme linked immunosorbent assay (ELISA)

The mouse colon cell line STC-1 (RRID: CVCL_J405) was purchased from American type culture collection (ATCC). The cells were cultured in DEME growth medium supplemented with 10% fetal bovine serum at 37°C and 5% CO_2_. Cells were plated into a 24-well plate with an appropriate density and then exposed to vehicle (DMSO, the control, less than 0.1%) with or without 80 μM of CDCA. After incubation, the culture supernatants were collected and the secreted GLP-1 was detected according to the manufacturers’ instructions by ELISA assay.

### 2.11 RNA isolation and quantitative reverse transcription PCR

Total RNA was isolated using 1 mL Trizol (Invitrogen). The cDNA templates were obtained from 500 ng of purified RNA using iScript Reverse Transcription Supermix for RT-qPCR (Bio-rad, CA). 1×SYBR Green Master Mix buffer (Takara, Otsu, Japan) was used for quantitative RT-PCR and assays were performed on a Roche lightCycler 480 II PCR machine. Gene specific primers were listed in **Table 1**. The targeted gene levels were normalized to β-actin housekeeping gene levels and the results were analyzed using the 2^-ΔΔCt^ method.

**Table 1.**
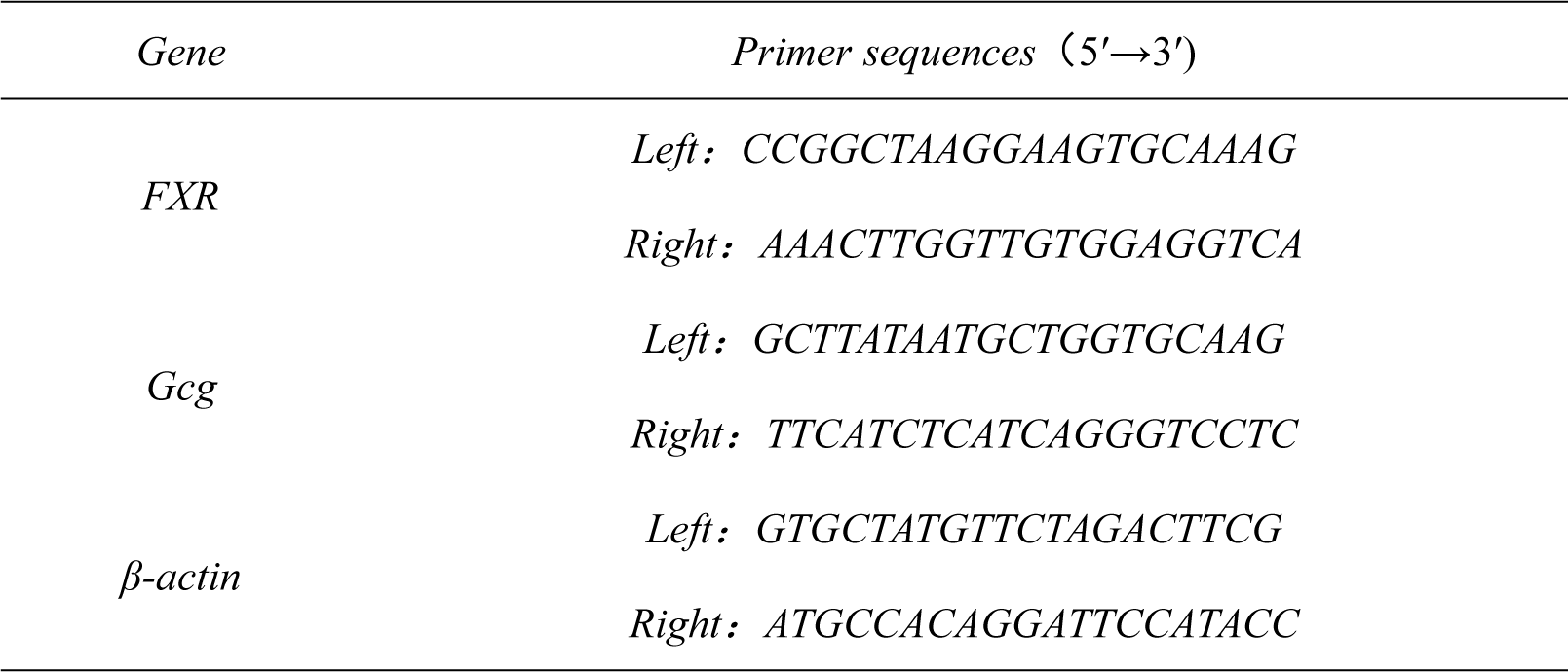
qRT-PCR primer sequences.

### 2.12 Statistical analysis

Results were presented as mean ± SD. Statistical significance between different groups was analyzed using the unpaired Student’s t-test for two-group comparison and one-way ANOVA followed by the Tukey-Kramer post hoc test. Statistical analyses were calculated using GraphPad prism 9.2 (GraphPad software, LaJolla, CA, USA). Correlations between BAs and microbiome abundances were performed using Spearman’s correlation analysis. Differences between experimental groups were considered significant at p < 0.05.

## 3. Results

### 3.1 DHM improves insulin resistance by the enhancement of incretin effect in HFD-induced mice

Mice were administrated with HFD with or without DHM for 8 weeks. There was no significant differences in food intake among different groups (Supplement data 1A and B). The body weights of mice were measured each week (Fig. 1A), and HFD administration led to a significant increase in the body weight at the end of the experiment, which was ameliorated by DHM intervention (Fig. 1B). To further investigate the effect of DHM on glucose homeostasis and insulin sensitivity, the OGTT and ITT assays were conducted. It was showed that DHM could significantly relieve glucose tolerance abnormalities caused by HFD (Fig. 1C and D). And ITT assay showed that HFD enhanced insulin resistance compared with the control group, while the changes were alleviated by DHM treatment (Fig. 1E and F). The incretin effect describes the phenomenon that oral glucose administration results in higher insulin secretion than intraperitoneal glucose administration (Kaur *et al*, 2015). To investigate the impact of DHM on incretin effect in HFD-induced mice, we performed IPGTT and OGTT experiments. Combined with the results of IPGTT and OGTT, it was showed that HFD could disrupt the incretin effect of mice, which was dominantly restored by DHM administration (Fig. 1G-J). Meanwhile, fasting blood glucose level was notably increased in HFD group, which was significantly inhibited by DHM treatment (Fig. 1K). In terms of serum lipid metabolism changes, the results showed that compared with the control group, the serum CHO (Fig. 1L), LDL-C (Fig. 1M) and TG (Fig. 1M) levels in HFD group were significantly increased, while HDL-C (Fig. 1O) level was significantly decreased (p<0.05). However, the changes of CHO, LDL-C, TC and HDL-C levels in the HFD group were effectively alleviated by DHM treatment, respectively (Fig. 1L-O). These results suggested that the administration of DHM could effectively improve the HFD-induced insulin resistance as well as abnormal glucose and lipid metabolism in mice, which might be related to the amelioration of the incretin effect.

**Figure 1.**
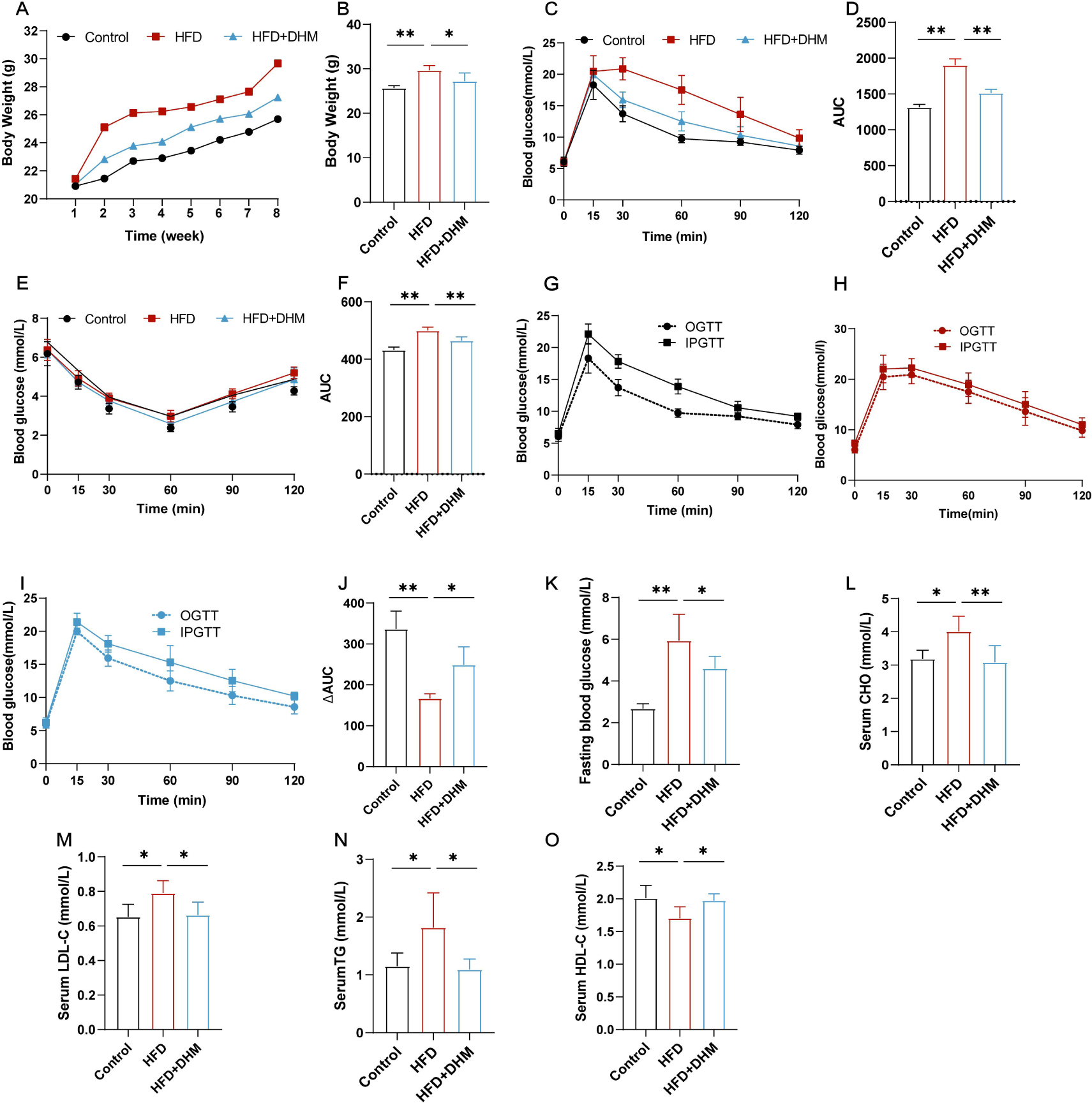
DHM improves insulin resistance by the amelioration of incretin effect in HFD-induced mice. (A) Body weights at each week among different groups. (B) Body weight at the end of experiment among different groups. (C) Oral glucose tolerance test (OGTT). (D) Area under curve (AUC) of OGTT. (E) Insulin resistance test (ITT). (F) Area under curve (AUC) of ITT. To determine the potential role of the effect in the influence of DHM on insulin secretion, mice in the control group (G), HFD (H), HFD+DHM (I) were subjected to the IPGTT and OGTT. (J) The ΔAUC is indicated by the shaded portions in G-J. (L) Fasting blood glucose at the end of experiment among different groups. (K) Serum CHO (L), LDL-C (M), TG (N), HDL-C (O) among different groups. All data reflect at least two independent experiments, and mean ± SD are plotted. *P < 0.05, **P < 0.01, results as demonstrated by one-way ANOVA followed by Tukey-Kramer post hoc test.

### 3.2 DHM increases serum GLP-1 level by improving GLP-1 secretion in intestine and inhibiting GLP-1 decomposition in HFD-induced mice

GLP-1 is a short endocrine peptide synthesized and secreted mainly by intestinal L cells (Hira *et al*, 2020). It helps prevent insulin resistance by promoting insulin production, lowering glucagon levels, inhibiting gastric emptying, reducing food intake, lowering blood sugar levels, and controlling weight(Mather, 2014). Therefore, we used immunofluorescence assay to investigate the expression of GLP-1 protein in the small intestine of mice (Fig. 2A). We found that GLP-1 protein expression was significantly decreased in HFD group compared with the control group, while the expression of GLP-1 was enhanced after DHM treatment (Fig. 2B). Meanwhile, DHM intervention increased serum GLP-1 level (Fig. 2C) and the relative expression of Gcg mRNA in intestinal tissue (Fig.2D). Taken together, the results showed DHM enhanced intestinal GLP-1 secretion and serum GLP-1 level in HFD-induced mice. Intestinal intraepithelial lymphocytes (IELs) are a kind of special T cell population in intestinal mucosal epithelium (Lockhart *et al*, 2024), and their different subsets and phenotypes are closely related to many diseases (Zhou *et al*, 2019). Thus, we further investigated whether DHM-induced change of serum GLP-1 level was resulted from the increase of its synthesis and secretion or the decrease of IELs’ degradation. We used flow cytometry to detect the expression of CD26 of IELs and the change of the proportion of each subset of IELs. The results were showed in Fig. 2E, F and Supplement data 2A. No statistically significant differences of IELs number were found among different groups (Fig. 2G). And there are no siginificant differences of TCRγδ^+^ and TCRαβ^+^ IELs among groups (Fig. 2H, I). For the analysis of TCRαβ^+^ IELs, the results showed that the proportion of TCRαβ^+^ CD8αα^+^ IELs showed no significant changes among the groups, while the proportion of TCRαβ^+^ CD8αβ^+^ IELs changed significantly (Fig. 2J, K). It showed HFD caused an increase in the proportion of TCRαβ^+^ CD8αβ^+^ IELs, which was decreased after DHM intervention. The CD26 was highly expressed in HFD group, which was suppressed after DHM intervention, suggesting the inhibitory effect on CD26 expression by DHM treatment (Fig. 2L). The proportion of CD26 secreted by TCRγδ^+^ IELs was no significant difference between HFD and HFD+DHM group (Fig. 2M). The proportion of CD26 secreted by TCRαβ^+^ IELs was significantly increased in HFD group, and decreased after DHM intervention (Fig. 2N). The proportion of CD26 secreted by TCRαβ^+^ CD8αβ^+^ IELs was increased in HFD, and decreased after DHM intervention (Fig. 2O), but no significant changes had found in the proportion of CD26 secreted by TCRαβ^+^ CD8αα^+^ IELs among the groups (Supplement data 2B). Therefore, we speculate that DHM intervention could increase GLP-1 secretion level in intestine of HFD-induced mice, and inhibit CD26 expression of IELs, particularly the expression of CD26 in TCRαβ^+^ CD8αβ^+^ IELs, finally increase serum GLP-1 level.

**Figure 2.**
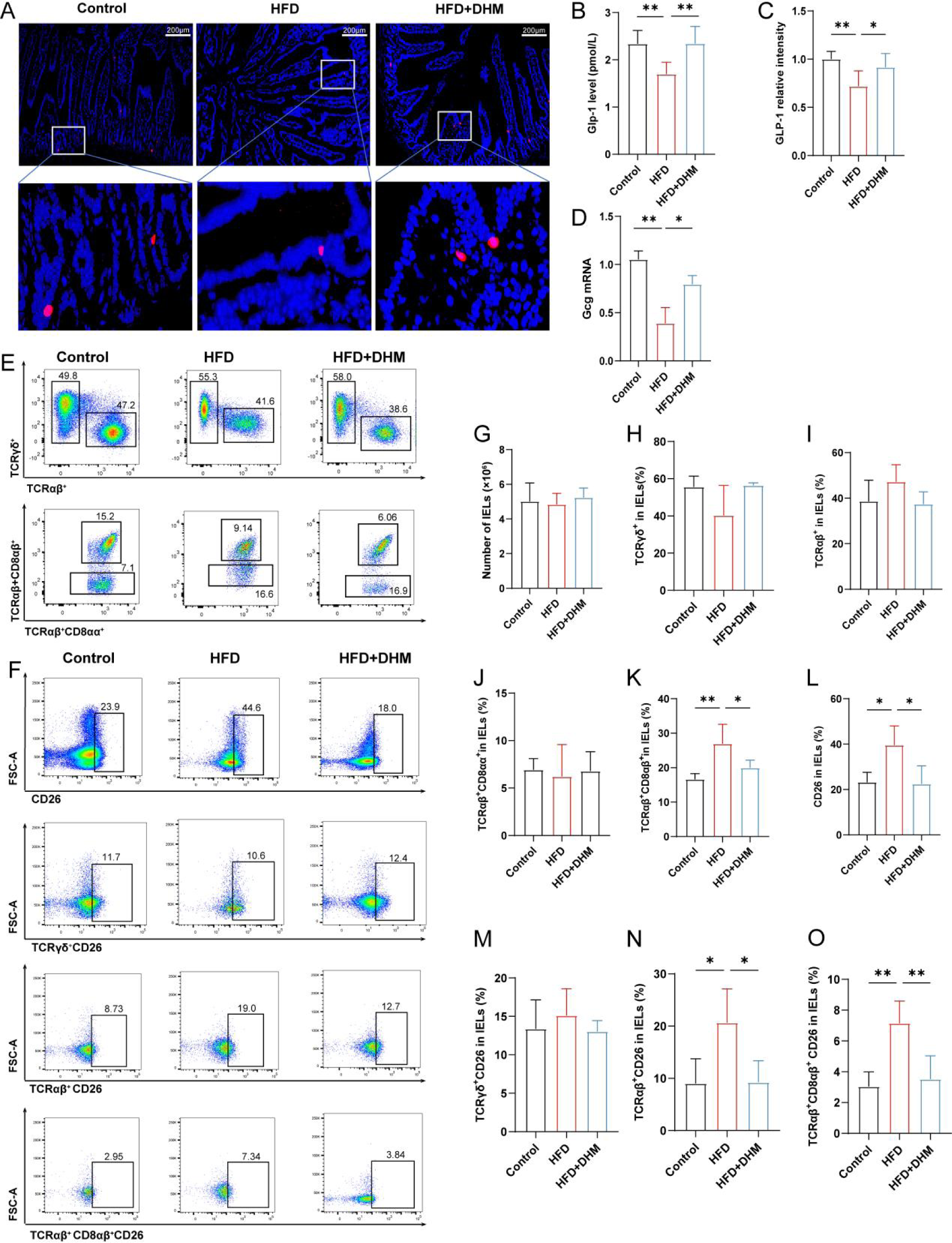
DHM increases serum GLP-1 level by enhancing GLP-1 secretion in intestine and inhibiting GLP-1 decomposition in HFD-induced mice. (A) Representative photograph of sections of small intestine stained with anti-GLP-1 antibody. Blue fluorescence indicates DAPI and red fluorescence indicates GLP-1 (n=3 per group). mice fed with Control, HFD, HFD+DHM over 8 week time. (B) Relative intensity of GLP-1 among groups (n=6). (C) The serum GLP-1 level among groups (n=6). (D) RT-qPCR measurements of the expression levels of Gcg mRNA in small intestine tissues of mice fed with different diets. (E) Flow fractionation of IELs subsets. (F) The expression of CD26 in IELs subsets. (G) Number of IELs in three groups. (H) The proportion of TCRγδ^+^ IELs in the indicated groups. (I) The proportion of TCRαβ^+^ IELs in the indicated groups. (J) The proportion of TCRαβ^+^CD8αα^+^ IELs in the indicated groups. (K) The proportion of TCRαβ^+^CD8αβ^+^ IELs in the indicated groups. (L) The expression of CD26 in IELs in the indicated groups. (M)The expression of CD26 in TCRγδ^+^ IELs in the indicated groups. (N) The expression of CD26 in TCRαβ^+^ IELs in the indicated groups. (O) The expression of CD26 in TCRαβ^+^ CD8αβ^+^ IELs in the indicated groups. All data represent at least two independent experiments. Mean ± SD are plotted. *P < 0.05, **P < 0.01 compared between the indicated groups results as demonstrated by one-way ANOVA followed by Tukey-Kramer post hoc test.

### 3.3 DHM ameliorates GLP-1 level and insulin resistance by modulation of gut microbiota in HFD-induced mice

In order to further verify the role of gut microbiota in the regulation of systemic insulin resistance by DHM, we carried out gut microbiota clearance and gut microbiota transplantation experiments, respectively. There was no significant differences in food intake between the HFD+Abx and HFD+DHM+Abx groups during the gut microbiota clearance experiment (Fig. 3A). In the gut microbiota clearance experiment, we measured the fasting blood glucose of mice at the end of the fourth week and carried out OGTT experiment. The results showed that the impaired glucose tolerance in HFD+DHM+Abx group was significantly relieved and the fasting blood glucose was significantly decreased, when compared with HFD+Abx group before gut microbiota clearance (Fig. 3B-D). The antibiotic mixture was then added to the drinking water of the mice for 4 weeks of gut microbiota clearance. Interestingly, when the gut microbiota was cleared, the effect of DHM on fasting blood glucose and glucose tolerance was disappeared (Fig. 3E-G). The analysis of IPGTT combined with OGTT showed that there was no significant difference between the two groups in incretin effect after the elimination of gut microbiota after gut microbiota clearance (Fig. 3H-J). The immunofluorescence assay results showed that there were no significant differences in the secretion of GLP-1, intestinal Gcg mRNA and serum GLP-1 content between HFD+Abx and HFD+DHM+Abx groups after gut microbiota clearance, suggesting gut microbiota plays a key role in the improvement of systemic insulin resistance by DHM (Fig. 3K-N).

**Figure 3.**
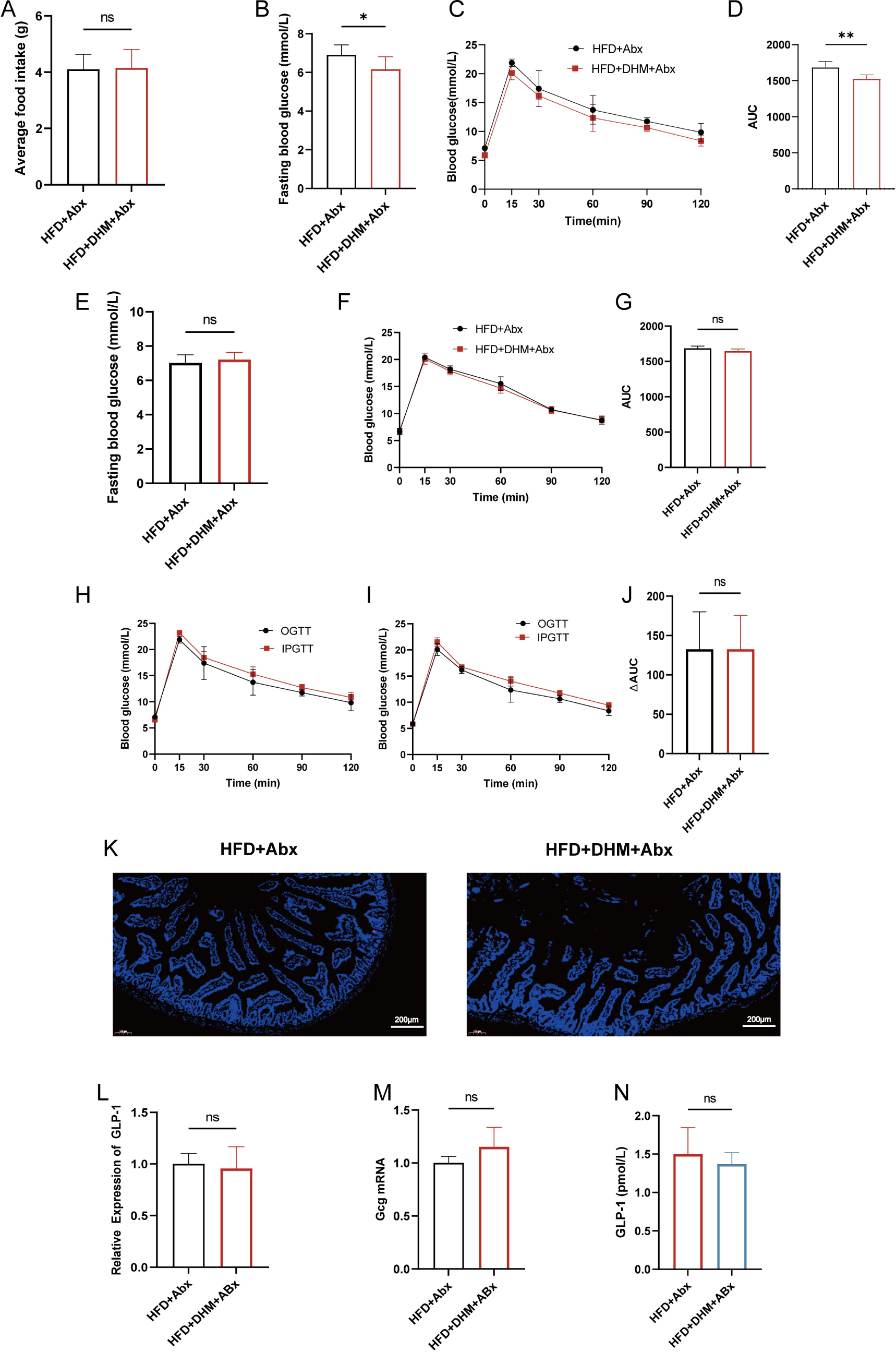
The improvement of DHM on fasting blood glucose, glucose tolerance, incretin and GLP-1 levels in mice induced by high-fat diet was significantly weakened after the intervention of antibiotics. (A) Average food intake. (B) Fasting blood glucose at the end of 4 week. (C) OGTT between HFD+Abx and HFD+DHM+Abx groups at the end of 4 week. (D) the area under the curve (AUC) in each group before gut microbiota clearance at the end of 4 week. (E) Fasting blood glucose after gut microbiota clearance at the end of 8 week. (F) OGTT between two groups after gut microbiota clearance at the end of 8 week. (G) the area under the curve (AUC) in each group after gut microbiota clearance at the end of 8 week. (H-J)To determine the potential role of the effect in the influence of DHM on insulin secretion, mice in HFD+Abx group (H), HFD+DHM+Abx group (I) were subjected to the IPGTT, and OGTT. (J) The ΔAUC is indicated by the shaded portions in H-I. (K) Representative photograph of sections of small intestine stained with anti-GLP-1 antibody. Blue fluorescence indicates DAPI and red fluorescence indicates GLP-1 (n=3 per group). mice fed with HFD+Abx and HFD+DHM+Abx over 8 week time. (L) Relative expression of GLP-1 in two groups (n=6). (M) RT-qPCR measurements of the expression levels of Gcg in small intestine tissues. (N) The GLP-1 level in two groups (n=9). All data reflect at least two independent experiments, and mean ± SD are plotted. *P < 0.05, **P < 0.01, ns, non-significance compared between the indicated groupsresults as demonstrated by unpaired Student’s t-test.

Furthermore, we carried out gut microbiota transplantation experiments, in order to further verify the role of gut microbiota in the regulation of insulin resistance. We performed gut microbiota clearance on mice during the 5^th^ and 6^th^ weeks of intervention and gut microbiota transplantation during the 7^th^ and 8^th^ weeks. Compared with Donor: HFD group and Donor: HFD+DHM group, there was no significant difference in food intake between the two groups during the experiment (Fig. 4A). The result of OGTT showed glucose tolerance impairment was relieved after gut microbiota transplantation (Fig. 4B-C). After gut microbiota transplantation, the improved effect of DHM on incretin effect was notably restored (Fig. 4D-F). The results of immunofluorescence and RT-qPCR assays showed that the intestinal expression of GLP-1 and Gcg mRNA, as well as serum GLP-1 were significantly increased (Fig. 4G-J). Taken together, these results suggest that the improvement of insulin resistance and GLP-1 level by DHM treatment may be mediated through the modulation of gut microbiota.

**Figure 4.**
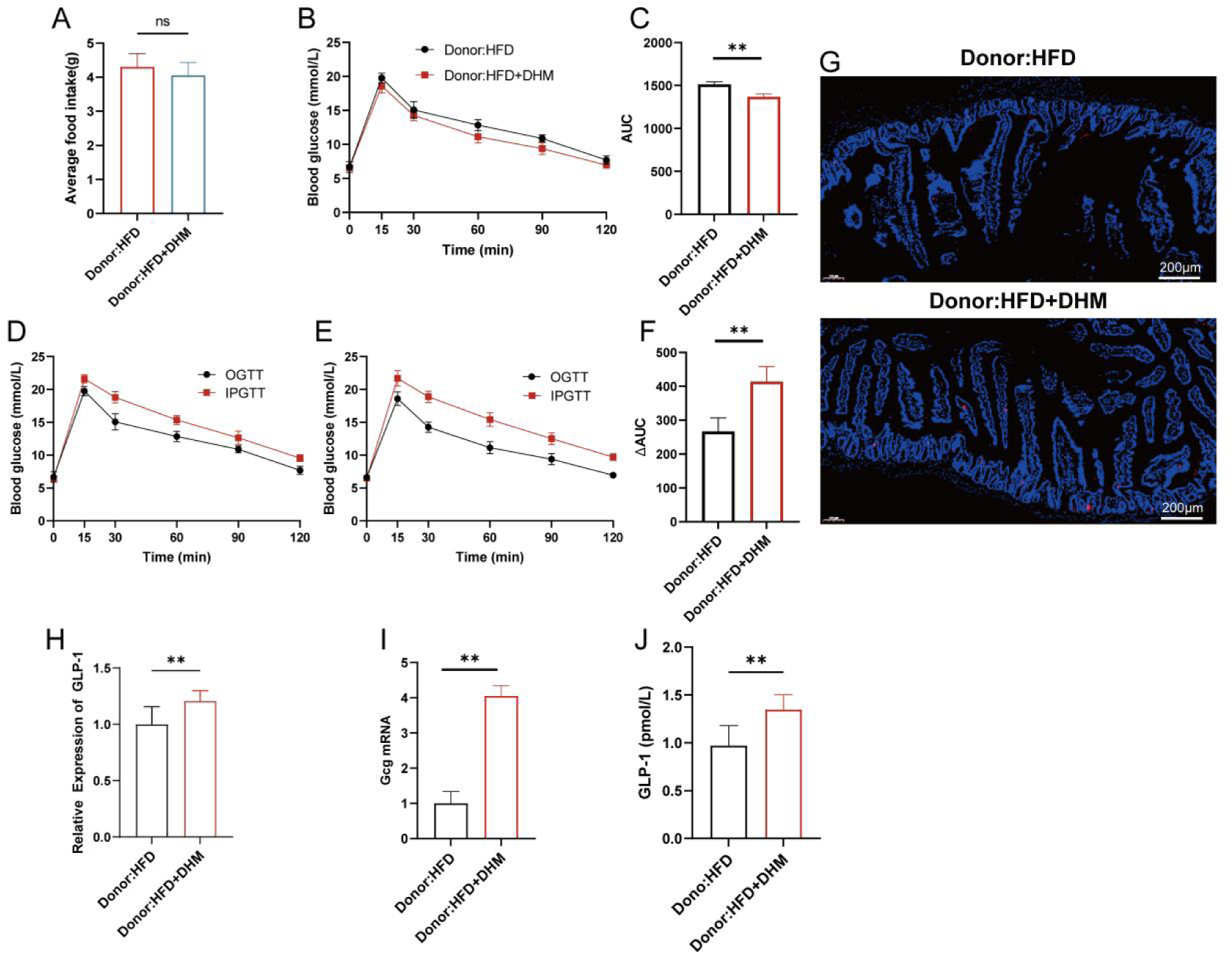
DHM could improve glucose tolerance, incretin, GLP-1 protein expression in intestinal tissue and serum GLP-1 level in mice induced by high-fat diet after gut microbiota transplantation. (A) Average food intake between Donor:HFD and Donor:HFD+DHM groups during 8 weeks; (B) OGTT between Donor:HFD and Donor:HFD+DHM groups. (C)the area under the curve (AUC) of OGTT in Donor:HFD and Donor:HFD+DHM groups. (D-F) To determine the potential role of the effect in the influence of gut microbial on insulin secretion in Donor:HFD group (D) and Donor:HFD+DHM group (E) were subjected to the IPGTT, and OGTT. (F) The ΔAUC is indicated by the shaded portions in R-S. (G) Representative photograph of sections of small intestine stained with anti-GLP-1 antibody.Blue fluorescence indicates DAPI and red fluorescence indicates GLP-1 (n=3 per group). Mice fed with Donor:HFD group and Donor:HFD+DHM group over 8 week time. (H) Relative intensity of GLP-1 in two groups(n=6). (I) RT-qPCR measurements of the expression levels of Gcg in small intestine tissues. (J) The GLP-1 level in two groups (n=9). All data reflect at least two independent experiments, and mean ± SD are plotted. *P < 0.05, **P < 0.01, ns, non-significance compared between the indicated groupsresults as demonstrated by unpaired Student’s t-test.

### 3.4 DHM changes the gut microbiota composition associated with insulin resistance in HFD-induced mice

To investigate how DHM modulates gut microbiota to improve GLP-1 secretion of intestinal L cells, we evaluated the gut microbiota biospectrum of gut microbiota by 16S rRNA sequencing and bioinformatics analysis. The results showed that compared with the control group, Ace, Chao, Sobs and Shannon indexes of HFD group were significantly different (p<0.05), while Sobs index was different between HFD and HFD+DHM group. And there was no significant difference among other three indexes (Fig. 5A). Meanwhile, PCoA test on OTU level showed that gut microbiota diversity distribution in HFD group was significantly different from that in the other two groups (Fig. 5B). Collectively, the gut microbiota diversity of mice in HFD group may be significantly changed.

**Figure 5.**
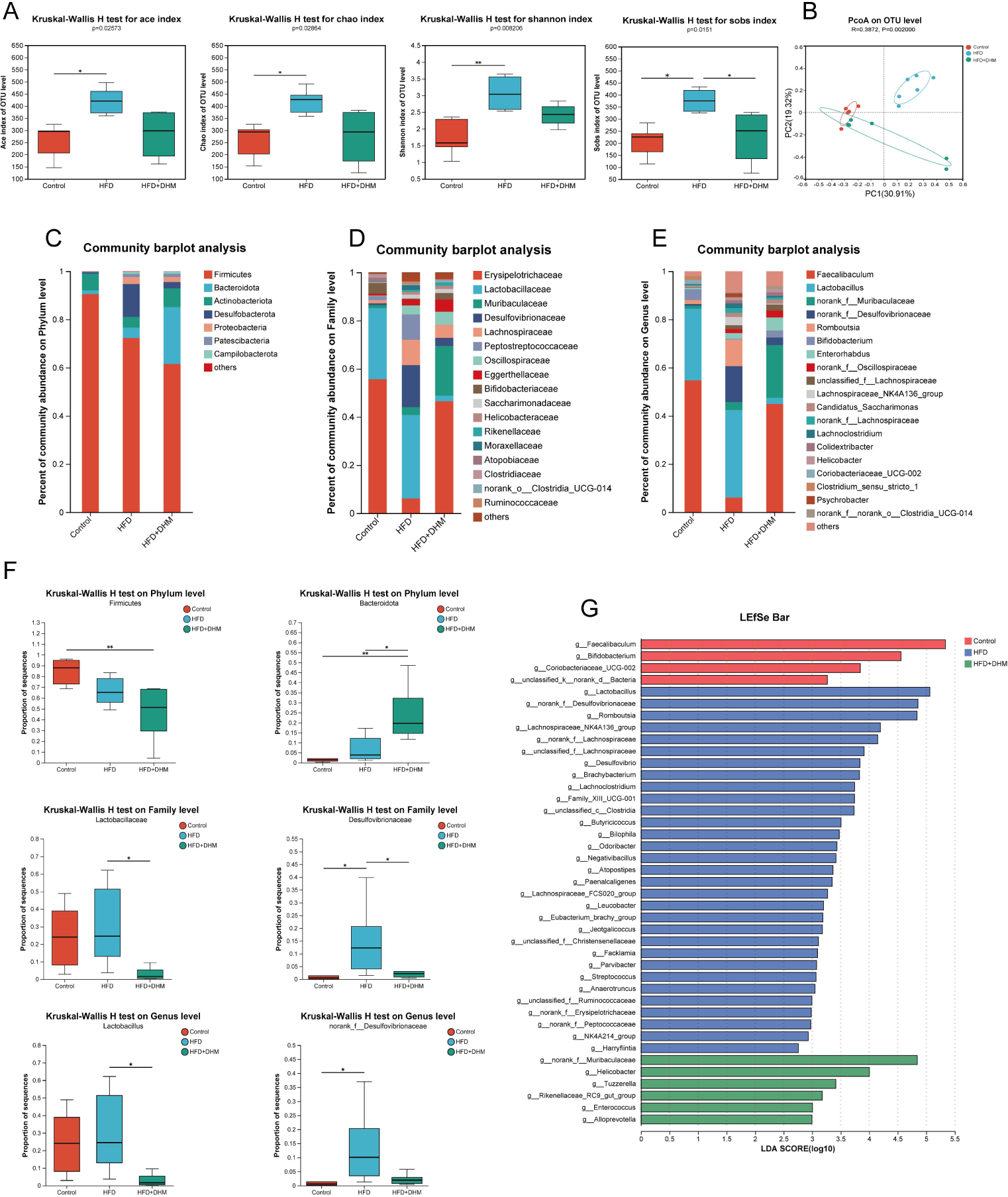
DHM changes the gut microbiota composition associated with insulin resistance in HFD-induced mice. (A) α diversity analysis. (B) PCoA analysis on OTU level among three groups. (C-E) Bar plot of gut microbiota composition on Phylum (C), Family (D), Genus level (E). (F) Relative proportion of sequence in the three groups (n=6 per group). (G) Taxa identified by LEfSe of samples from the three groups (LDA≥2 and p≤0.05). Mean± SD are plotted. * P < 0.05, **P < 0.01 as determined by Kruskal-Wallis H test.

To explore the relationship between DHM and changes of the gut microbiota, we performed a classification analysis of OTUs from primary sequencing data, revealing changes in microbiota composition of the indicated three groups. The results showed that the gut microbiota was changed obviously on Phylum, Family and Genus levels (Fig. 5 C-E). On phylum level, the abundance of *Bacteroidata* and *Firmicutes* had no significantly differences between the control group and HFD group, whereas the abundance of *Bacteroidata* was increased in HFD+DHM group (Fig. 5F). The changes between HFD and HFD+DHM was similar to the result between the control group and HFD group (Fig. 5F); however, DHM significantly decreased the abundance of *Firmicutes* in HFD+DHM group and DHM group compared with the control group. On the Family level, compared with the control group, the abundance of *Desulfovvibrionaceae* in HFD group was increased significantly, which was decreased notably after DHM intervention. The abundance of *Lactobacillaceae* was changed significantly in HFD+DHM group when compared with HFD group, and has no signicifcantly differences between the control group and HFD group (Fig. 4F). On Genus level, *norank_f_Desulfovibrionaceae’s* abundance was increased in HFD group; the abundance of *Lactobacillus* had a tendency to increase after HFD, but there was no significant difference, and it was decreased significantly after DHM intervention (Fig. 5F). To identify specific gut microbiota associated with HFD-induced insulin resistance, we examined the composition of gut microbiota among the three groups by LEfSe. The results showed that *Lactobacillus* and *norank_f_Desulfovibrionaceae* were enriched in HFD group, while *norank_f_Muribaculaceae* were enriched in HFD+DHM group (Fig. 5G). These results suggest that DHM can change the gut microbiota composition associated with insulin resistance in HFD-induced mice.

In order to analyze the relationship between differential gut microbials and insulin resistance index, we carried out Spearman correlation analysis of relevant parameters. The results of Spearman correlation analysis showed that the increase of *Desulfobacterota* abundance was positively correlated with GLu, TG, CHO and body weight, and negatively correlated with GLP-1. DHM changed the gut microbiota structure of insulin resistance mice induced by HFD (Supplement data 3A-C).

The results of 16S rRNA sequencing of gut microbiota showed that DHM could significantly improve the increase of *Lactobacillaceae* abundance in mice induced by HFD. It has been reported that *Lactobacillaceae* can participate in the regulation of bile acid metabolism by encoding bile salt hydrolase (BSH). The results of genus-level difference analysis showed that DHM could effectively inhibit the increase of *Lactobacillus* induced by HFD. In order to determine whether DHM has a regulatory effect on the BSH activity of mice induced by HFD, we measured and evaluated the BSH activity of mice in each group (Supplement data 4). The results showed that the BSH activity in HFD group was significantly higher than that of Control group, and the BSH activity in HFD+DHM group was significantly lower than that of HFD group. This result was similar to the change trend of *Lactobacillaceae* abundance. Therefore, it is speculated that DHM can reshape the gut microbiota structure and regulate BSH activity in HFD-induced mice.

### 3.5 DHM could modulate intestinal microbiota metabolic pathways in HFD-induced mice

In order to verify the effect of DHM on intestinal metabolites in HFD group, we collected feces from each group of mice at the 8^th^ week, and performed non-targeted metabolomics analysis on the collected fecal samples by LC/MC. The results of PLS-DA showed that the metabolites of each group were well-separated from HFD group, and the results of permutation test showed the prediction ability of the PLS-DA model was good and there was no over-fitting phenomenon. (Fig. 6A-D).

**Figure 6.**
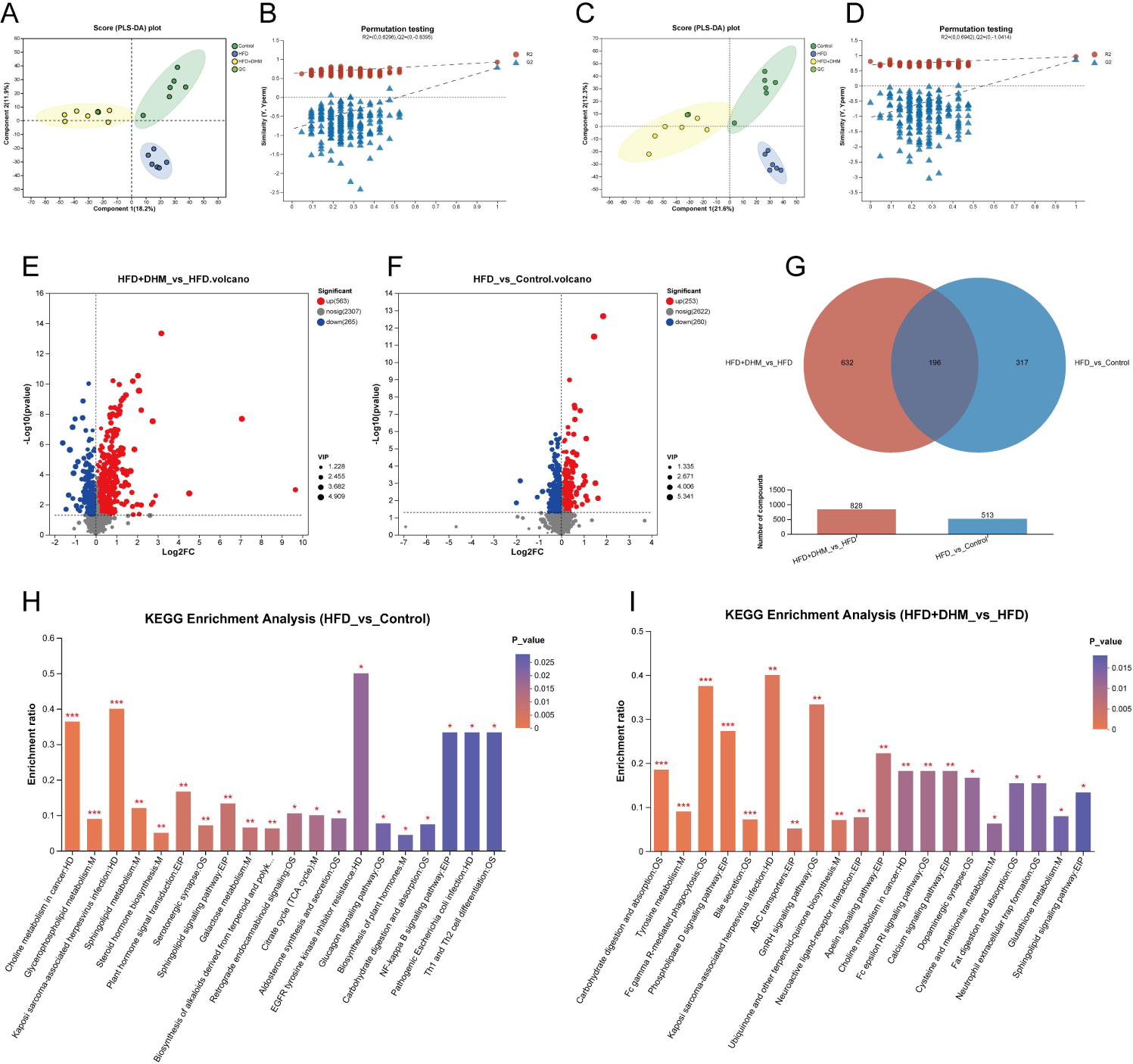
DHM can regulate intestinal metabolism pathways of mice induced by HFD. (A) the postive modes’ PLS-DA score plots of fecal samples from three groups. (B) Permutation tests of the three groups. (C) the negative modes’ PLS-DA score plots of fecal samples from threer groups. (D) Permutation tests of the three groups. (E-F) Volcano plot of fecal metabolomics of fecal samples showing the significantliy modes in HFD vs. Control (E) and HFD+DHM vs. HFD (F). (G) Venn diagram of fecal samples between HFD+DHM vs HFD and HFD vs Control. (H-I) KEGG annotation analysis of the changed metabolites based the differences.

To screen for differential metabolites among the indicated three groups, we set a threshold of VIP>1.0,FC>1.0 or <1, P<0.05. In the comparison between the HFD and the control groups, 253 differential metabolites were up-regulated, while 260 were down-regulated (Fig. 6E). In the comparison between the HFD+DHM group and the HFD group, 563 differential metabolites were up-regulated and 265 down-regulated after DHM intervention (Fig. 6F). In addition, there were 196 common differential metabolites between the two groups (Fig. 6G).

In order to analyze the effects of differential metabolites, we used the KEGG database to conduct pathway enrichment analysis of the measured metabolic sets, and identified the most important biochemical metabolic pathways and signal transduction pathways involved in differential metabolites (Fig. 6H and I). There were 43 important differential pathways (P<0.05) between HFD group and the control group, and 39 differential metabolic pathways (P<0.05) between HFD+DHM group and HFD group. To further analyze the differential pathways, we used the corrected P<0.05 to screen the differential pathways and the differential metabolite pathways involved were counted (Table 2), between HFD+DHM and HFD group, there were 5 different pathways, including Fc gamma R-mediated phagocytosis, Phospholipase D signaling pathway, Tyrosine metabolism pathway, Carbohydrate digestion and absorption pathway and Bile secretion pathway. These results suggest that DHM could regulate the intestinal microbiota metabolic pathways of mice induced by HFD, and bile acid metabolism might be highly involved with the improvement of insulin resistance by DHM treatment.

**Table 2.**
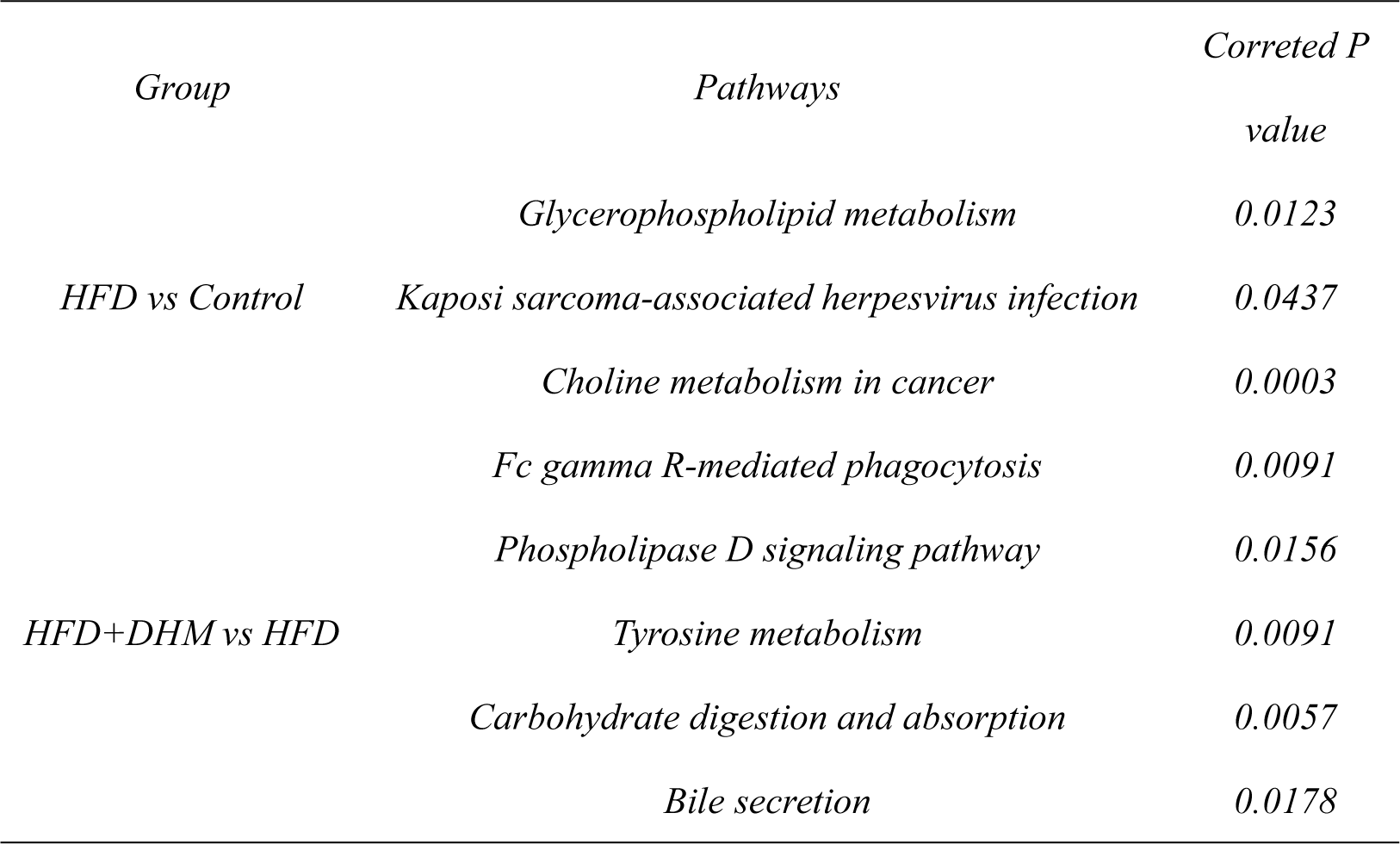
Significant Enrichement Pathways in HFD vs Control and HFD+DHM vs HFD.

### 3.6 DHM regulates intestinal bile acid metabolism and inhibits CDCA and FXR expression in HFD-induced mice

Bile acids exerts several functions on regulating intestinal metabolism(Cai *et al*, 2022). To explore the role of bile acid metabolism in the improvement of insulin resistance by DHM, we collected mouse feces and conducted a targeted analysis of bile acid metabolism. PLS-DA analysis showed that the main components of fecal bile acids in mice of HFD group were significantly separated from the other three groups, and the main components of HFD+DHM were significantly overlapped with those of the control group (Fig. 7A). It was suggested that the bile acid metabolism in HFD mice was abnormally regulated, and DHM intervention could significantly alleviate this change, which may be consistent with that in the control group. In order to analyze the changes of intestinal bile acids in mice, we further detected the composition of intestinal bile acids in mice. Conjugated and unconjugated bile acids were counted in each group (Fig. 7B and C). The results showed that the level of total bile acid in HFD group was significantly higher than that of the control group (Fig. 7E). Similar results can also be observed when analyzing conjugated and unconjugated bile acids (Fig. 7D). However, we found that there was no significant change in the conjugated/unconjugated bile acid ratio between HFD and the control groups. However, DHM significantly reduced the conjugated/unconjugated bile acid ratio between HFD+DHM and HFD groups (Fig. 7F). It was suggested that DHM could significantly change the intestinal bile acid metabolism in HFD group. Next, we further analyzed the changes of primary and secondary bile acids, and found that HFD significantly increased the content of primary bile acids in the intestine, and DHM significantly decreased the content of primary bile acids, while there was no significant change in secondary bile acids (Fig. 7G). Meanwhile, the primary/secondary bile acid ratio showed DHM could effectively alleviate the increase of primary bile acids (Fig. 7H). This suggests that DHM could correct abnormal primary bile acid metabolism induced by HFD, but had no significantly effect on secondary bile acids. When analyzing the bile acids with metabolic regulation ability, it showed the ratio of FXR-activated bile acids to FXR-inhibited bile acids was significantly increased in HFD compared with the control group, while it was significantly decreased after DHM intervention (Fig. 7I). FXR agonistic bile acids included Cholic acid (CA), Taurodeoxycholic acid (TCA), Tauro-β-muricholic acid (T-β-MCA), Tauroursodeoxycholic acid (TUDCA) and CDCA. It showed that CDCA was significantly increased in HFD group, which was significantly inhibited by DHM treatment. There were no significant changes in other FXR agonistic bile acids among groups (Fig. 7J-M). CDCA is reported to be the natural ligand of FXR, so we tested the FXR expression in each group. The result showed the expression of FXR mRNA and protein were significantly increased in HFD group and decreased after DHM, respectively (Fig. 7N-O).

**Figure 7.**
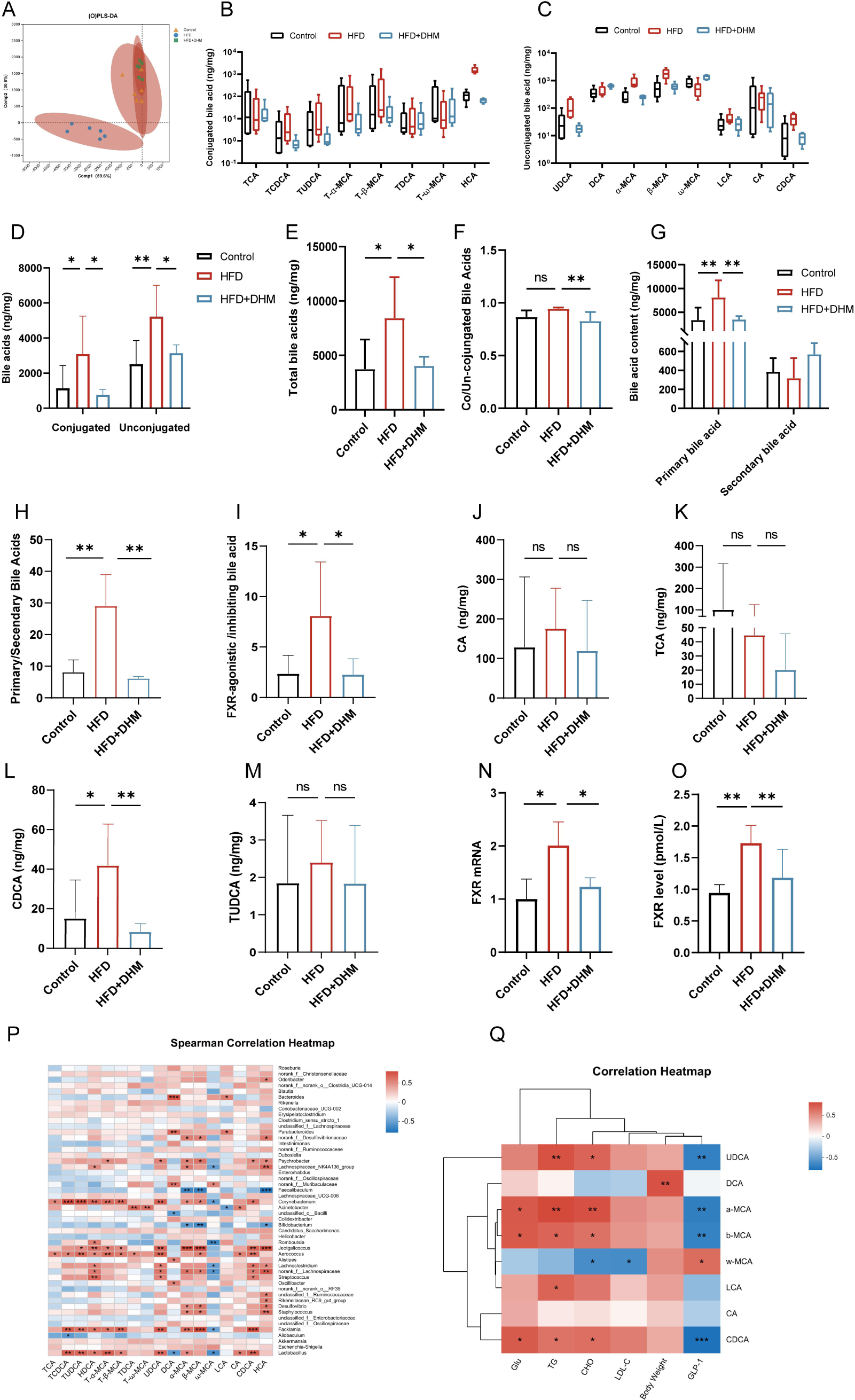
DHM regulates intestinal bile acid metabolism in mice induced by HFD. (A) PLS-DA score plots of fecal samples from three groups. (B-C) Statistics of conjugated and unconjugated bile acids in mouse feces. (D) Differences in conjugated and unconjugated bile acids among the three groups. (E) Difference of total bile acid among groups. (F) Statistics of the ratio of conjugated bile acid to unconjugated bile acid in each group. (G) Contents of primary and secondary bile acids in each group. (H) Ratio of primary bile acid to secondary bile acid in each group.(I)Ratio of FXR receptor agonistic and inhibitory bile acid abundance.(J-M) Levels of CA, TCA, CDCA, TUDCA in mouse fecal samples. (N) Relative Gcg mRNA expression levels in three groups. (O) The content of FXR in the small intestine of three groups of mice. (P) Sperman Correlation Heapmap on Genus level between gut microbial and bile acids. (Q) Sperman Correlation Heapmap between bile acids and normal indexs.All data reflect at least two independent experiments, and mean ± SD are plotted.*P < 0.05, **P < 0.01, ***P < 0.0001, ns, non-significance results as demonstrated by one-way ANOVA followed by Tukey-Kramer post hoc test

In order to further clarify the relationship between gut microbiota and bile acid metabolic profiles, the correlation between gut microbiota at genus level and bile acid profiles was analyzed. As shown in Figure 7P, the contents of CDCA, UDCA and TUDCA showed a significant positive correlation with *Lactobacillaceae*. There was a significant positive correlation between *Desulfovibrio* and CDCA, MCA, DCA and LCA. In order to further expound the relationship between body mass, serum indexes, serum GLP-1 levels and bile acid profiles, the correlation analysis was carried out between them and bile acid profile. As shown in Fig. 7Q, CDCA was positively correlated with body weight, blood glucose, TG, CHO and LDL-C, and negatively correlated with serum GLP-1 level. There was a significant positive correlation between α-MCA and TG, CHO, and a significant negative correlation between α-MCA and GLP-1. There was a significant positive correlation between β-MCA and blood glucose and a significant negative correlation between β-MCA and GLP-1, respectively.

### 3.7 CDCA inhibits GLP-1 secretion by stimulating and increasing FXR expression in intestinal L cells

In order to further verify the effect of CDCA on GLP-1 secretion by intestinal L cells, we next used STC-1 cells to further investigate how CDCA inhibits GLP-1 secretion. STC-1 cells were stimulated with different concentrations of DHM (1, 5, 10, 20, 40, 60, 80 and 100 μM) for 24 h, and then we assessed cell viability by the CCK-8 assay and measuring GLP-1 levels in the harvested culture supernatant at 0, 10, 40, 80 μM. Our results showed that at different concentrations, CDCA had no significant effect on the proliferation of STC-1 cells during the range from 1μM to 80 μM. At 100 μM, the viability of STC-1 cells was significantly inhibited (Fig. 8A). However, compared with the control group, 80 µM of CDCA significantly inhibited the secretion of GLP-1 by STC-1 cells (Fig. 8B-D). STC-1 cells were further treated with 80 µM of CDCA in the presence of absence of FXR antagonist Z-Gug (10 μM) for 24 h. The results showed that CDCA resulted in an increased expression of FXR mRNA, decreased expression of Gcg mRNA and the secretion of GLP-1, respectively. In the presence of Z-Gug, the expression of FXR mRNA was decreased significantly, and the expression of Gcg mRNA and the secretion of GLP-1 were increased, respectively (Fig. 8E-H). To sum up, CDCA promotes the relative mRNA expression of FXR and inhibits the relative mRNA expression of Gcg and the expression of GLP-1 in intestinal L cells.

**Figure 8.**
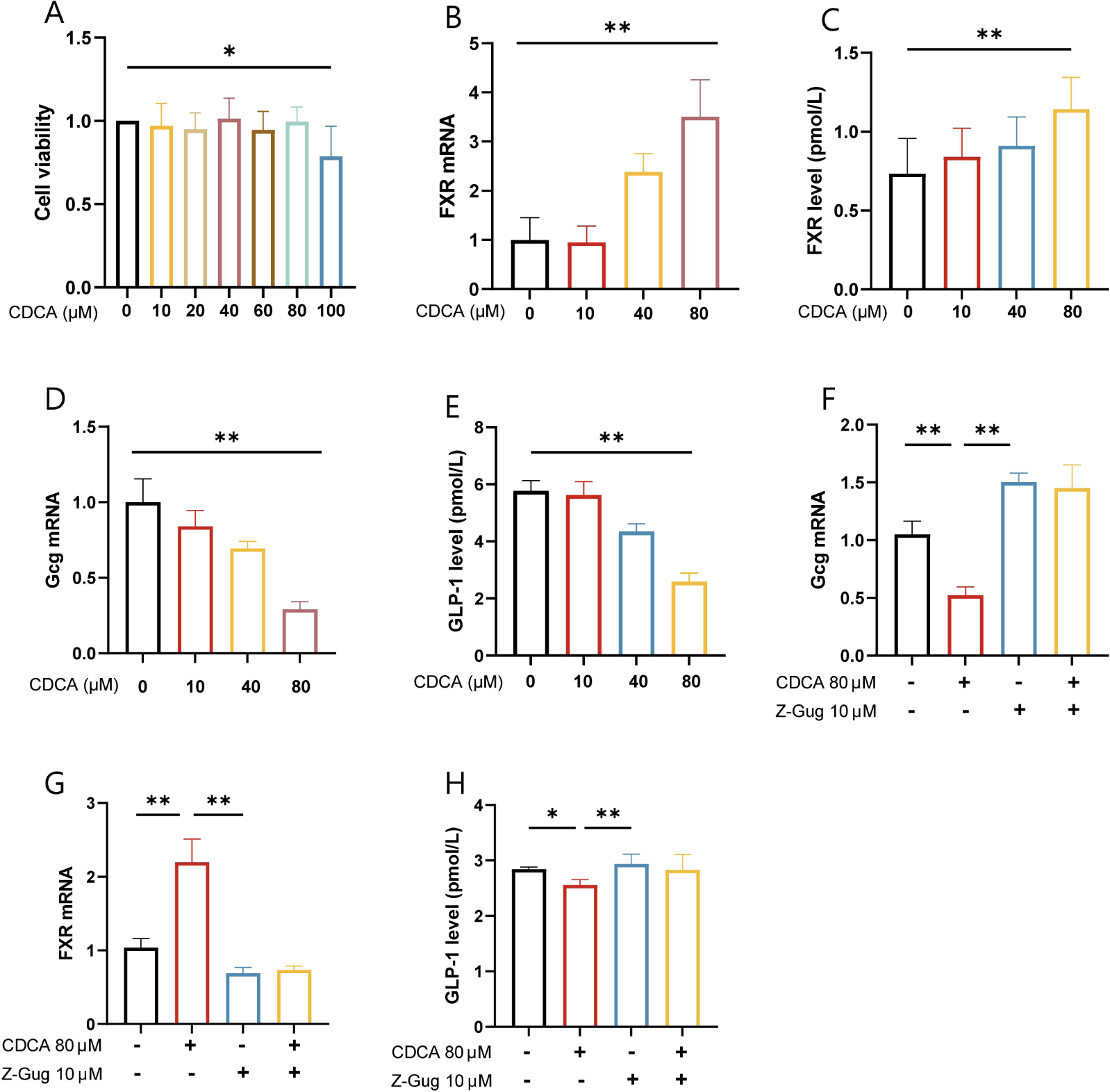
CDCA inhibits GLP-1 secretion by stimulating and increasing FXR expression in intestinal L cells. (A) Cell viability analysis with 0-100 μM CDCA. (B-C) Relative mRNA expression levels of FXR,Gcg in four groups with 0-80 μM CDCA. (D) GLP-1 content secreted by STC-1 cell with 0-80 μM CDCA. (E-G)With or without 80 μM CDCA and 10 μM Z-Gug, STC-1 secreted GLP-1 and relative expression of Gcg, FXR. All data reflect at least two independent experiments, and mean ± SD are plotted.*P < 0.05, **P < 0.01, ***P < 0.0001, ns, non-significance results as demonstrated by one-way ANOVA followed by Tukey-Kramer post hoc test.

## 4. Discussion

Insulin resistance has been considered as a common risk factor for a variety of metabolic diseases, which can lead to the occurrence and development of a variety of metabolic diseases (Padilla *et al*, 2022). The pathogenesis of insulin resistance is complex. Gut microbiota (York, 2023; Takeuchi *et al*, 2023) and metabolite disorder, excessive energy intake and intestinal immunity (Malesza *et al*, 2021) play vital roles in the pathogenesis of insulin resistance. At present, there is no effective new drug treatment for insulin resistance (Fahed *et al*, 2022). DHM is mostly extracted from a woody vine of the genus *Ampelopsis* of the family Vitaceae, and is also extracted from Hovenia dulcis Thunb (Sun *et al*, 2022). DHM has a potential role in the prevention and treatment of a variety of chronic metabolic diseases (Wang *et al*, 2023b). A large number of experiments have proved that DHM can reduce atherosclerosis, improve blood glucose and blood lipids (Hou *et al*, 2021; Janilkarn-Urena *et al*, 2023). Glucagon-like peptide-1 (GLP-1) secreted by intestinal L cells, which can regulate blood glucose and insulin resistance. GLP-1 is rapidly broken down by DPP4/CD26 (Müller *et al*, 2019), which is predominantly expressed on the surface of intestinal intraepithelial lymphocytes (IELs). In this study,we found that DHM could promote the synthesis and secretion of GLP-1 in intestinal L cells through the “gut microbiota-CDCA” pathway; meanwhile, it could regulate the proportion of TCRαβ^+^ CD8αβ^+^ IELs and the expression of CD26, reduce the degradation of GLP-1, thus increasing serum GLP-1 level and alleviating insulin resistance (Fig. 9).

**Figure 9.**
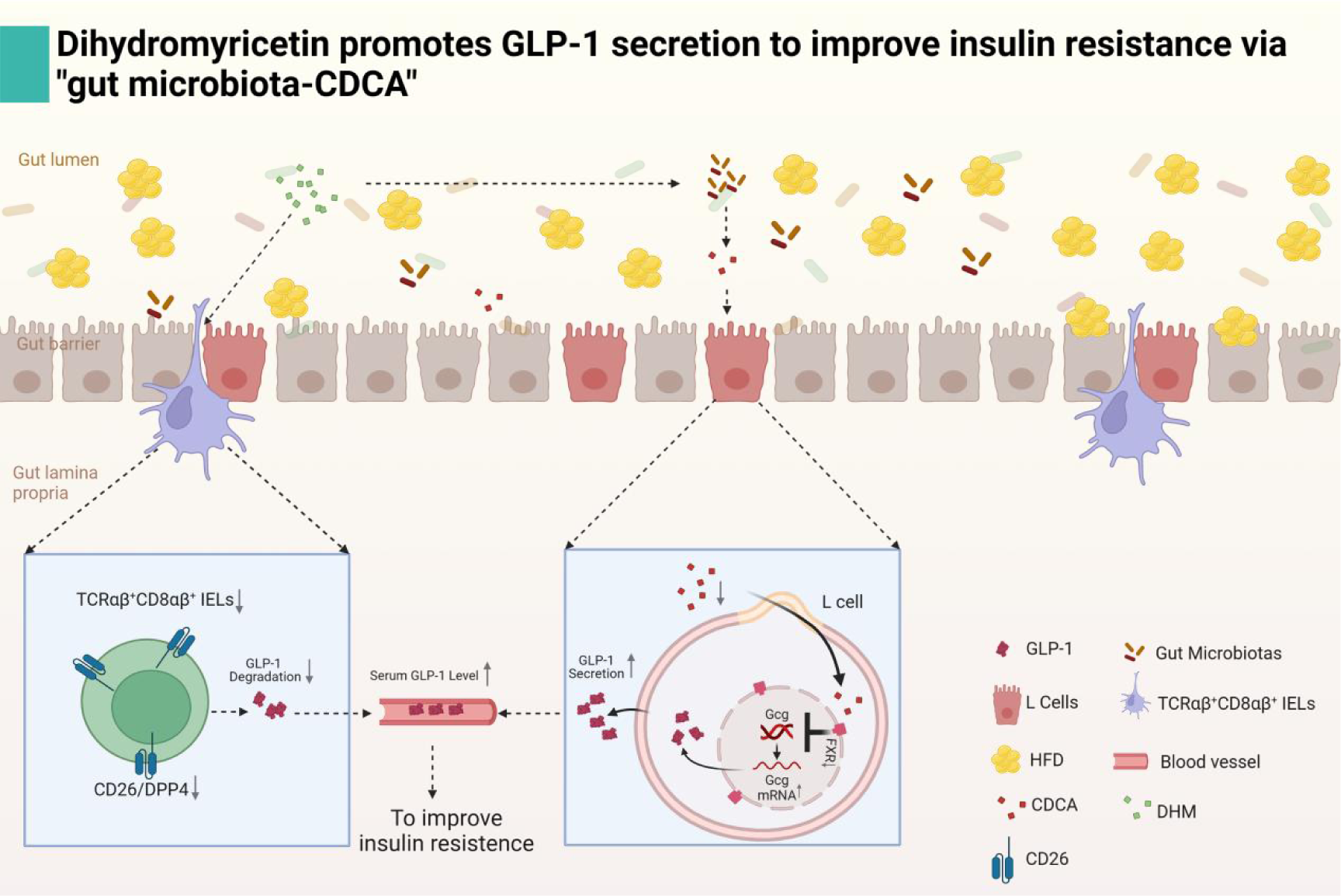
Dihydromyricetin promotes GLP-1 secretion to improve insulin resistance via “gut microbiota-CDCA”. DHM could promote the synthesis and secretion of GLP-1 in intestinal L cells through the “gut microbiota-CDCA” pathway. Meanwhile, it could regulate the proportion of IELs and the expression of CD26 in IELs and TCRαβ^+^ CD8αβ^+^ IELs, reduce the degradation of GLP-1, resulting in an increased serum GLP-1 level and ameliorated insulin resistance.

Firstly, we established a HFD-induced insulin resistance model. Then, we investigated the effects of DHM on glucose metabolism and insulin resistance in mice. The results showed that DHM could significantly reduce the serum glucose, and significantly alleviate insulin resistance and impaired glucose tolerance, which are the core index for evaluating diabetes. In addition, DHM also affected the lipid metabolism of HFD mice. DHM could effectively alleviate the changes of TG, CHO, LDL and HDL in HFD group. The incretin effect describes the phenomenon that oral glucose leads to higher insulin secretion than intraperitoneal glucose. The incretin effect refers to the effect of oral glucose that results in higher insulin secretion than intraperitoneal glucose. To determine a potential role in the effects of DHM on incretin effect, we performed IPGTT and OGTT experiments. The results of OGTT combined with IPGTT showed that the incretin effect of HFD mice was significantly weaker than that of normal mice based on isoglycemic conditions; after DHM intervention, the incertin effect of mice in HFD+DHM group was enhanced significantly than that of HFD mice. By detecting the level of GLP-1 in the serum, we found that the serum GLP-1 level in HFD group was decreased significantly, and increased after DHM intervention. Immunofluorescence showed that the expression of GLP-1 in the intestinal tissue of HFD mice was significantly increased after DHM treatment. The expression of Gcg mRNA was consistent with the result of immunofluorescence.

Dipeptidyl peptidase 4 (DPP-4/CD26), a serine protease belonging to the type II transmembrane glycoprotein family, is expressed on the surface of T cells, NK cells, B cells (Hu *et al*, 2022). It has enzymatic activity and can selectively scavenge N-terminal dipeptides of polypeptides and proteins containing proline or alanine at the second position. GLP-1 secreted by intestinal L cells is rapidly degraded by CD26/DPP4, limiting its biological function (Sun *et al*, 2020). IELs are a unique T cell population located in the intestinal mucosa, which closely contacts and interacts with intestinal epithelial cells. It has been found that integrin β7^+^ IELs are involved in the development of insulin resistance. Intestinal L cells in β7^-/-^ mice are significantly increased, GLP-1 levels are increased, and glucose tolerance is significantly improved. When β7^-/-^ mice are fed with high-fat and high-sugar diet, they have significant resistance to obesity, diabetes and so on(He *et al*, 2019). This study suggests that some special subsets of intestinal IELs may affect the expression level of gut-derived GLP-1 and regulate the metabolism of the body, so interventions targeting IELs may become a new strategy to regulate the level of GLP-1. In order to verify whether IELs was involved in the regulation of GLP-1 physiological concentration maintenance, we separated IELs and detected them by flow cytometry. The results showed that there was no significant change in the number of IELs in each group, but the expression of CD26 in HFD group was increased and decreased after DHM intervention. The expression of CD26 was decreased after DHM intervention. TCRαβ^+^CD8αβ^+^ IELs were significantly increased in HFD, but the proportion decreased after DHM intervention. CD26 secreted by TCRαβ^+^CD8αβ^+^ IELs was enhanced in HFD group, but was decreased after DHM intervention. It was suggested that TCRαβ^+^CD8αβ^+^ IELs were involved in the regulation of GLP-1 secretion.

It has been reported in many literatures that the gut microbiota plays an important role in the regulation of body metabolism (Fan & Pedersen, 2021; Winston & Theriot, 2020). However, whether gut microbiota plays an important role in the regulation of intestinal GLP-1 secretion by DHM is unknown. In order to verify whether the gut microbiota plays a vital role in the secretion of GLP-1, we carried out gut microbiota clearance and transplantation experiments. The experimental results are the same as expected. There were significant glucose intolerance and insulin resistance between HFD+DHM+Abx group and HFD+Abx group before the elimination of gut microbiota. After the elimination of gut microbiota, the difference of insulin resistance between the two groups was inhibited, and GLP-1 secretion was no different change. In the experiment of gut microbiota transplantation, GLP-1 secretion inhibition, insulin resistance and impaired glucose tolerance were observed to be alleviated after gut microbiota transplantation. The results suggested that the gut microbiota played a positive regulatory role in the secretion of GLP-1.

Gut microbiota, as an important parasitic microorganism in the host intestine, fully participates in the nutritional metabolism of the host and is mainly affected by diet (Fan & Pedersen, 2021; Puljiz *et al*, 2023). Obesity induced by HFD has been shown to be associated with gut microbiota dysregulation, as demonstrated by a reduction in gut microbiota diversity and an alteration in gut microbiome composition (Geng *et al*, 2022; Van Hul & Cani, 2023). Therefore, we measured the gut microbiota of mice to further study its regulatory mechanism. The results of 16S rRNA analysis showed that the diversity and abundance of gut microbial in HFD group changed significantly. DHM could modulate gut microbiota diversity, especially an decrease of *Desulfovvibrionaceae*, revealing protective features. At the genus level, DHM can effectively regulate the abundance changes of *norank_f_Desulfovibrionaceae* and *Lactobacillus* caused by HFD. The regulation of these key microorganisms by DHM is closely related to their therapeutic effects on insulin resistance, which was confirmed by correlation analysis.

Accumulating evidence supports the important role of gut microbiota composition in influencing host metabolism, and suggests that targeting gut microbiota composition can regulate host metabolism and reverse insulin resistance (Schoeler & Caesar, 2019). The effect of gut microbiota on host energy metabolism is mainly through amino acids, vitamins, short-chain fatty acids (Xiao *et al*, 2022; Colombo *et al*), bile acids (Winston & Theriot, 2020) and other metabolites, which have changed to varying degrees in obesity. Metabolite profiles of HFD-induced mice and DHM treated mice were analyzed by non-targeted metabonomics. In this study, we show that long-term HFD interferes with physiological and biochemical processes in mice by affecting clusters of relevant metabolic pathways. Metabolite patterns in HFD-induced mice were significantly different from those in healthy mice, depending on alterations in glycerophospholipid metabolism and choline metabolism, and changes in FcγR-mediated phagocytosis. Meanwhile, phospholipase D signaling pathway, tyrosine metabolism, carbohydrate digestion and absorption, and bile acid metabolism are also closely related to the intervention of DHM on HFD-induced obesity (Table 2).

Bile acids are important physiological agents for the absorption of intestinal nutrients and the secretion of lipids, toxic metabolites and xenobiotics by the biliary tract (Agus *et al*, 2021). In the intestinal lumen, different bile acids have different physiological functions (Collins *et al*, 2023; Fogelson *et al*, 2023). Meanwhile, bile acids are also important signaling molecules and metabolic regulators, which can activate nuclear receptors to regulate glucose and energy homeostasis and maintain metabolic homeostasis (Chiang & Ferrell, 2020). So, we performed a targeted metabolic assay for bile acid metabolism of mice. The results showed that the intestinal bile acid metabolism of HFD mice changed significantly. Total bile acids and primary bile acids were significantly increased. Further analysis of bile acids with biological utility showed that FXR receptor agonistic bile acids increased significantly in HFD group and decreased significantly after DHM intervention. Subsequently, we analyzed several FXR-agonistic bile acid concentration changes and found that CDCA was significantly increased in HFD and decreased after DHM intervention. It had been found that FXR receptor could participate in the regulation of GLP-1 secretion in intestinal cells. The results of ELISA and RT-qPCR showed that the expression of FXR was enhanced in HFD group and decreased after DHM intervention. It has been reported that CDCA is a natural agonist of FXR, which can activate FXR to participate in a variety of metabolic processes in the intestine. Then we detected the expression of FXR mRNA and protein in intestinal tissue by TR-qPCR and ELISA.

The results showed that the expression of FXR mRNA and protein in intestinal tissue in high-fat diet group were increased, and decreased after DHM intervention. At the same time, the content of BSH was detected by ELISA. It was found that the content of intestinal BSH in mice was increased by high-fat diet, and the content was decreased by DHM intervention. In the analysis of the correlation between bile acid and gut microbiota, it was observed that the increase of *Lactobacillus* was positively correlated with CDCA. The major source of BSH in the intestine is *Lactobacilli*, which decompose conjugated bile acids into unconjugated bile acids. We hypothesized that the high-fat diet caused an increase in *Lactobacilli* and thus increased BSH content, which in turn increased CDCA synthesis in the intestines.

STC-1 cells are a commonly used intestinal L cell line in vitro (Zhu *et al*, 2022). To explore the effect of CDCA on GLP-1 secretion by intestinal L cells, we conducted an in vitro study using STC-1 cells. In STC-1 cells, CDCA significantly increased the contents of FXR and decreased the contents of Gcg mRNA and GLP-1. CDCA had no significant effect on cell viability at 0-80 μM. We treated STC-1 cells with FXR antagonists Z-Gug and CDCA, and the results showed that the expression of FXR receptor was increased, while the expression and secretion of Gcg mRNA and GLP-1 were decreased in the presence of CDCA alone; after FXR receptor was antagonized, the expression of Gcg mRNA and GLP-1 was increased.The results suggest that CDCA inhibits Gcg expression and reduces GLP-1 secretion by activating FXR.

This study has several limitations. First of all, we have not performed transplantation experiments with specific gut microbiota in mice to verify the role of differential gut microbiota in insulin resistance. Second, intestinal CDCA was found to be changed in HFD group, but no relevant animal supplementation experiments has been carried out. It is necessary to further improve the research to fully interpret the scientific hypothesis.

## ACKNOWLEDGEMENTS

This research was supported by the National Natural Science Foundation of China (No. 82173505) and Natural Science Foundation of Chongqing, China (cstc2020jcyj-msxmX0502)

## AUTHOR CONTRIBUTIONS

Long Yi conceived and designed the study. Mantian Mi provided guidance for the design of the experiment. Pengfei Li, and Yong zhang performed the experiments; Yu yao, Pengfei Hou and Ruiliang Zhang collected the samples. Pengfei Li analyzed the data and wrote the manuscript. Hedong Lang provided technical support, Qianyong Zhang and Xiaolian Wang participated in data analysis and checking. All authors contributed to the article and approved the submitted version.

## Abbreviations

Abx: antibiotic
AUC: area under curve
ATCC: American type culture collection
BSH: bile salt hydrolase
CA: cholic acid
CDCA: chenodeoxycholic acid
CHO: cholesterol
CCK-8: cell counting kit-8
DHM: dihydromyricetin
DPP-IV: ipeptidyl peptidase-4
ELISA: enzyme linked immunosorbent assay
FXR: farnesoid X receptor
GLP-1: glucagon-like peptide-1
Glu: glucose
Gcg: glucagon gene
HFD: high-fat diet
HDL-C: high-density lipoproteins
IELs: intestinal intraepithelial lymphocytes
IECs: intestinal epithelial cells
IPGTT: intraperitoneal glucose tolerance test
ITT: insulin tolerance test
LDL-C: low-density lipoproteins
OGTT: oral glucose tolerance test
OTU: operational taxonomic units
PCA: Principal Component Analysis
PBS: phosphate buffered saline
QC: quality control
qPCR: quantitative polymerase chain reaction
SPF: secific pathogen free
T-β-MCA: tauro-β-muricholic acid
TCA: taurochenodeoxycholic acid
TG: triacylglycerol
TUDCA: tauroursodeoxycholic acid
T2DM: diabetes mellitus type 2
Z-Gug: Z-Guggulsterone

---

**Supplementary data 1.**
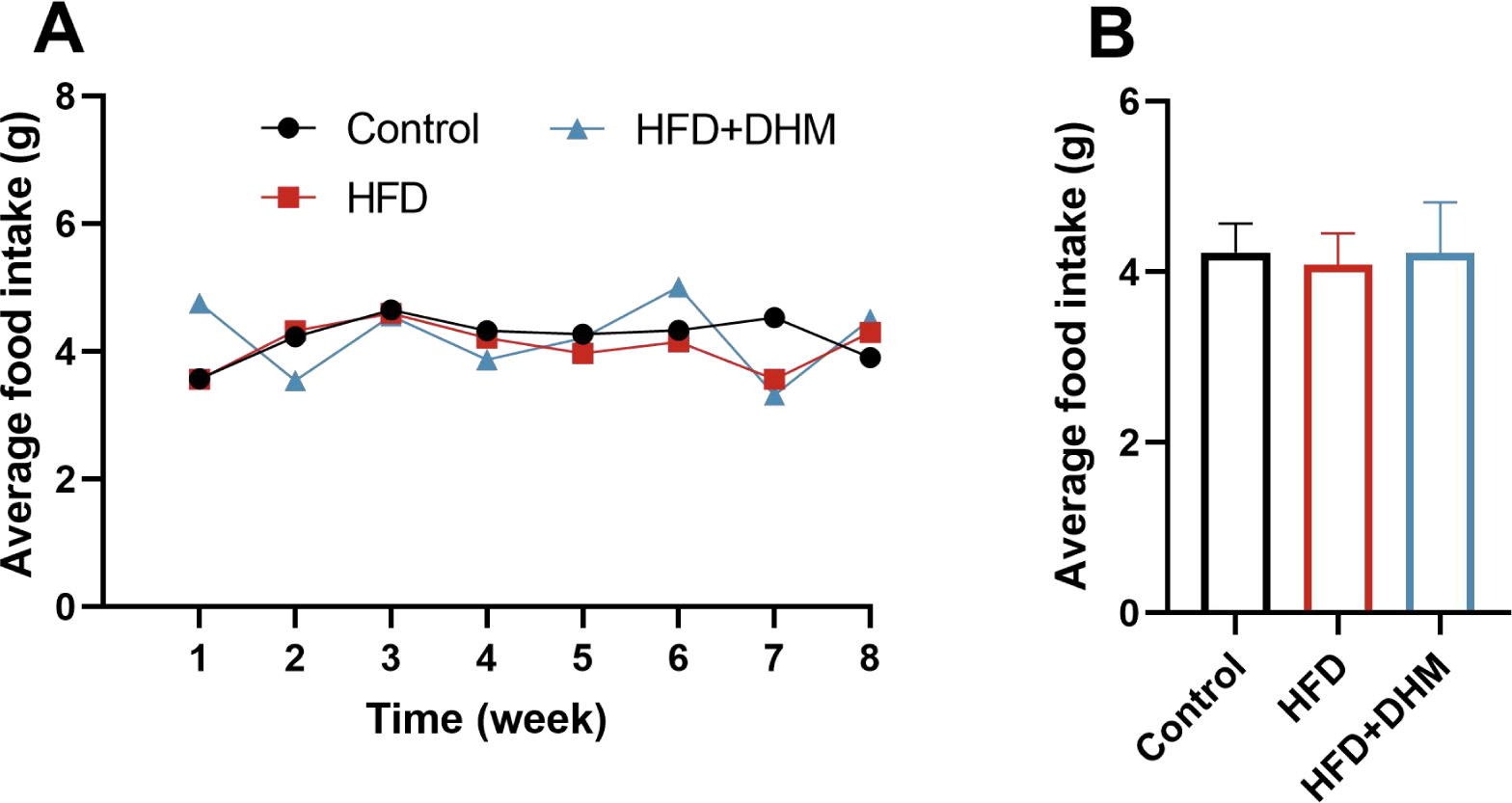
Food intake among different groups.

**Supplementary data 2.**
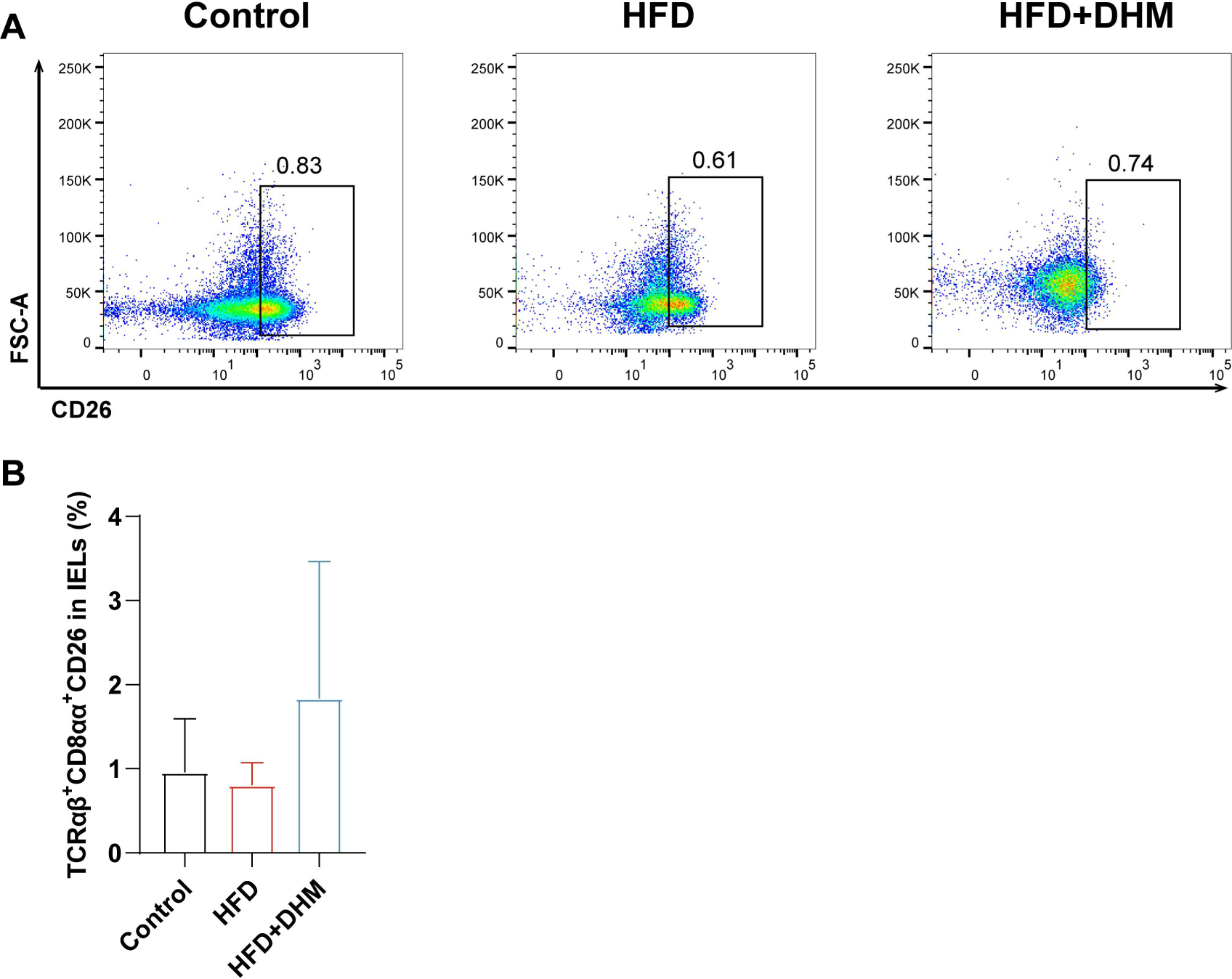
The proportion of TCRαβ^+^CD8αα^+^ IELs and the expression of CD26 in TCRαβ^+^CD8αα^+^ IELs.

**Supplementary data 3.**
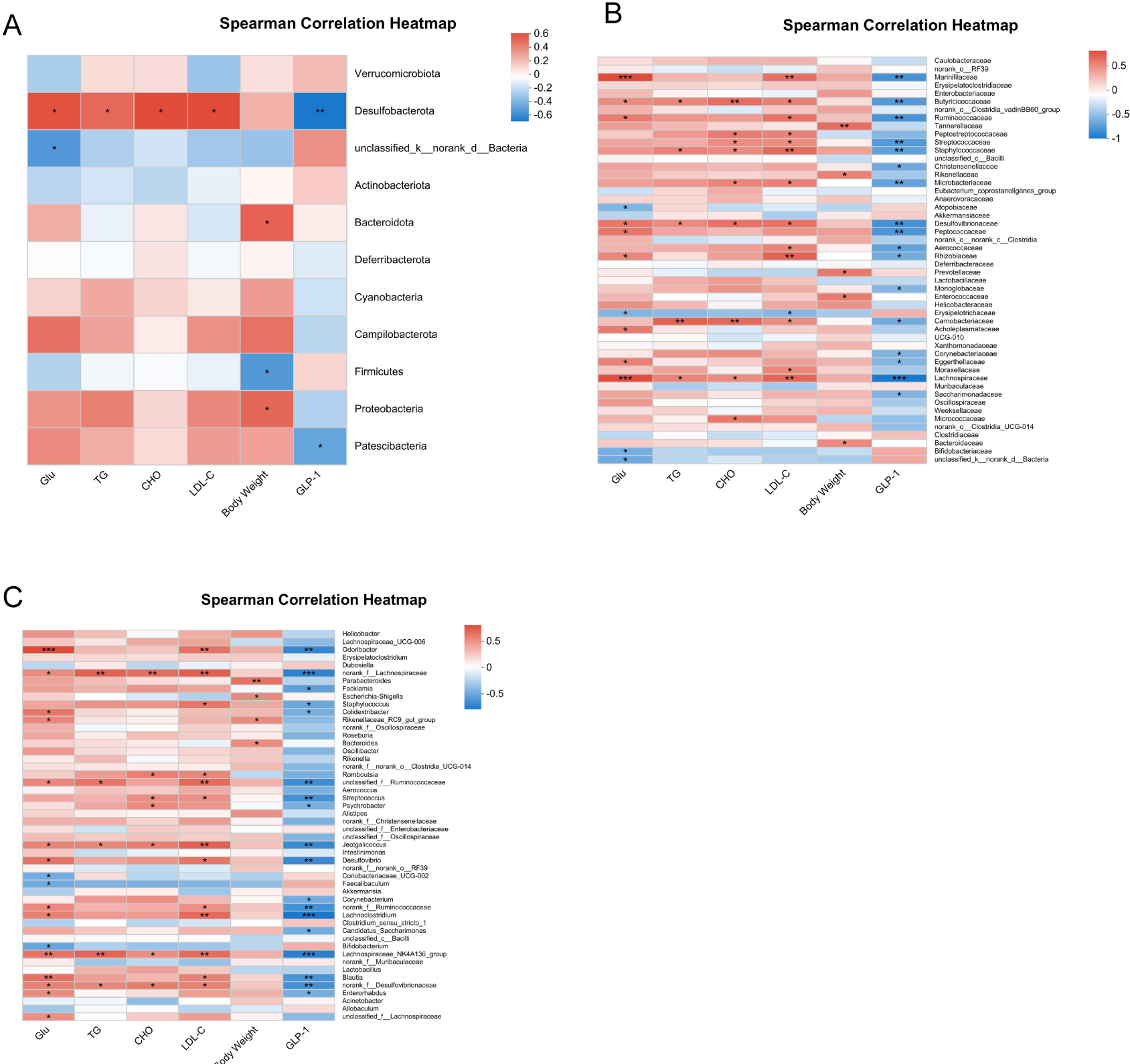
Spearman Correlation Heapmap analysis between gut microbial and general indicators. (A) Spearman Correlation Heapmap on Phylum level. (B) Spearman Correlattion Heapmap on Family level. (C) Spearman Corelation Heapmap on Genus level.

**Supplementary data 4.**
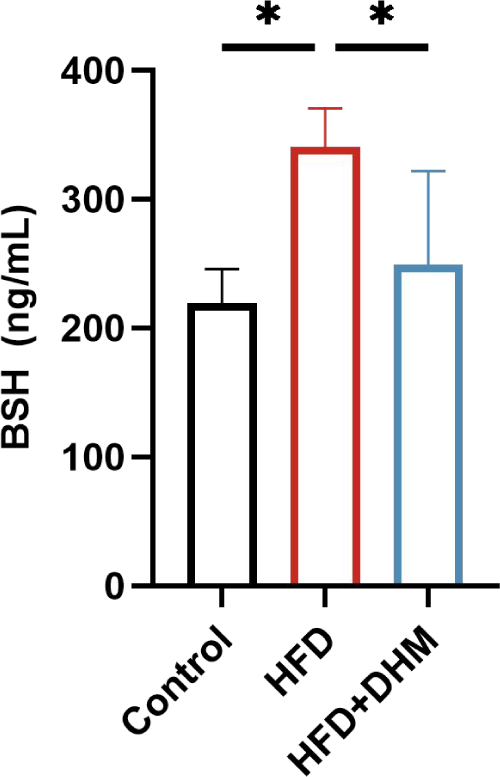
Content of intestinal bile salt hydrolase in mice.

